# Charting neuroimaging-based head circumference and cranial shape across development

**DOI:** 10.64898/2026.08.29.747918

**Authors:** P. Mattisson, A.S. Mandal, M. Gardner, A. Zapaishchykova, B. Jung, S. Prem, S. Karandikar, H.E. Akouri, D. Zimmerman, E. Levitis, J.A. Berken, G. Ball, R.A.I. Bethlehem, J.D. Bernstock, T.P. Roberts, L. Bloy, H. Huang, A. Vossough, S. Sotardi, E. Liao, M. Fisher, D.M. McDonald-McGinn, R.T. Shinohara, B.H. Kann, A.F. Alexander-Bloch, J. Seidlitz

**Affiliations:** Lifespan Brain Institute, The Children’s Hospital of Philadelphia and Penn Medicine, Philadelphia, PA, USA; Department of Psychiatry, University of Pennsylvania, Philadelphia, PA, USA; Department of Child and Adolescent Psychiatry and Behavioral Science, The Children’s Hospital of Philadelphia, Philadelphia, PA, USA; Department of Neurology, Mass General Brigham, Harvard Medical School, Boston, MA, USA; Artificial Intelligence in Medicine (AIM) Program, Mass General Brigham, Harvard Medical School, Boston, MA, USA; Department of Radiation Oncology, Dana-Farber Cancer Institute, Brigham and Women’s Hospital, Boston Children’s Hospital, Harvard Medical School, Boston, MA, USA; Perelman School of Medicine, University of Pennsylvania, Philadelphia, PA, USA; Division of Neonatology, The Children’s Hospital of Philadelphia, Philadelphia, Pennsylvania; Division of Gastroenterology, Hepatology, and Nutrition, The Children’s Hospital of Philadelphia, Philadelphia, Pennsylvania; Developmental Imaging, Murdoch Children’s Research Institute, Melbourne, Victoria, Australia; Department of Paediatrics, University of Melbourne, Victoria, Australia; Department of Psychology, University of Cambridge, Cambridge, UK; Division of Neurosurgery, Texas Children’s Hospital, Baylor College of Medicine, Houston, TX, USA; Program in Advanced Imaging Research, Department of Radiology, The Children’s Hospital of Philadelphia, Philadelphia, PA, USA; Department of Radiology, University of Pennsylvania, Philadelphia, PA, USA; Division of Plastic and Reconstructive Surgery, Children’s Hospital of Philadelphia, Philadelphia, Pennsylvania; Department of Surgery, University of Pennsylvania, Philadelphia, Pennsylvania; Department of Oncology, Perelman School of Medicine, University of Pennsylvania, Philadelphia, PA, USA; Division of Oncology, Children’s Hospital of Philadelphia, PA, USA; 22q and You Center, Children’s Hospital of Philadelphia, Philadelphia, Pennsylvania; Division of Genetic and Genomic Medicine, Children’s Hospital of Philadelphia, Philadelphia, Pennsylvania; Department of Pediatrics, University of Pennsylvania, Philadelphia, Pennsylvania; Penn Statistics in Imaging and Visualization Center, Department of Biostatistics, Epidemiology, and Informatics, Perelman School of Medicine, University of Pennsylvania, Philadelphia, Pennsylvania; Center for Artificial Intelligence and Data Science for Integrated Diagnostics (Ai2D), Perelman School of Medicine, University of Pennsylvania, Philadelphia, Pennsylvania; Institute for Translational Medicine and Therapeutics, University of Pennsylvania, Philadelphia, PA, USA

## Abstract

Quantitative assessment of head growth supports early detection of neurological disease, yet current clinical practice relies on manual measurements that are variable, anatomically limited, and difficult to scale. Here we introduce a fully automated imaging-native framework (CranioTrace) that transforms routine neuroimaging into standardized, population-referenced markers of cranial development. By integrating topology-constrained segmentation, contour regularization, and geometry-based quality control for automated slice selection, CranioTrace robustly extracts head circumference and cranial morphology from heterogeneous MRI and CT data without manual intervention. The framework enabled construction of sex-specific population growth models across pediatric development in a dataset comprising 9,685 scans spanning birth to early adulthood across 25 cohorts. Imaging-derived head circumference shows strong agreement with tape measurements across independent datasets, high reproducibility (intraclass correlation up to 0.99), and consistent performance across modalities. Beyond conventional head circumference, CranioTrace quantifies cranial shape and asymmetry, capturing developmental dynamics that are not accessible through conventional measurements. Population modeling reveals rapid early-life expansion followed by nonlinear deceleration and stable sex differences. Application to neurogenetic cohorts identifies disease-consistent shifts in growth trajectories: head circumference was higher in neurofibromatosis type 1 and 16p11.2 deletion and lower in 22q11.2 deletion and 16p11.2 duplication. By converting clinical imaging archives into scalable cranial phenotypes linked to probabilistic reference models, CranioTrace provides a foundation for imaging-based growth charting, integrated skull– brain phenotyping, and precise assessment of neurodevelopment.

## Introduction

Quantifying head circumference and cranial shape from brain imaging offers a precise, reproducible, and clinically scalable complement to traditional physical exam methods, with growing potential to advance pediatric neurodiagnostics. While head circumference via external tape measure remains a cornerstone of early pediatric assessment – routinely used to monitor growth and flag neurological abnormalities^1,2^ – tape measurements are subject to high inter-assessor variability^3^ and may be difficult to obtain in children who are uncooperative, sedated, or have craniofacial anomalies. For retrospective studies, these measurements may also be missing from electronic health records^4^, difficult to link to hospital imaging data, or separated in time from subsequent neuroimaging. These physical measurements also provide limited insight into regional cranial morphology or internal brain structure, reducing their utility in detecting subtle or syndromic deviations in growth.

The addition of imaging-derived head circumference to clinical workflows offers an anatomically grounded measure that can be automatically extracted for those patients who have clinically indicated neuroimaging and serve as a complement to traditional tape measurements. Using volumetric scans such as MRI or CT, head circumference can be precisely calculated based on the true outer cranial boundary, ensuring high consistency across patients and institutions^5^.

These measurements can be derived retrospectively from clinical imaging archives to enable longitudinal comparisons. Importantly, advances in deep learning and image processing now make it feasible to deploy a single computational pipeline that operates across imaging modalities – including both MRI and CT – allowing for broad applicability in diverse clinical contexts, including emergency settings where CT is more commonly used and routine growth measurements are often not documented^6–9^.

Beyond measuring size, this approach also enables detailed assessment of cranial shape, which can reveal diagnostic insights into conditions such as craniosynostosis, hydrocephalus, or genetic syndromes with characteristic morphologies^10^. Shape-based features, extracted alongside head circumference, provide a richer phenotypic profile that can improve diagnostic specificity and support early detection of abnormal developmental trajectories. For example, assessing anterior–posterior elongation, occipital flattening, or asymmetry in skull shape can aid in differentiating normal variants from pathologic cranial deformities^11^.

When contextualized within population growth models, imaging-based head circumference and shape measurements become powerful tools for precision pediatric care. Similar to physical growth charts for height and weight, neuroimaging-based growth charts allow for age- and sex-adjusted centile scoring of cranial metrics^12,13^. These centiles can help detect conditions such as microcephaly, megalencephaly, or ventriculomegaly, even when outward physical signs are minimal or obscured by overlapping clinical features. They are particularly useful in complex neurogenetic or metabolic conditions, where early identification of abnormal growth patterns may guide further genetic testing, surveillance, or intervention^14,15^.

Integrating head circumference and shape analysis into neuroimaging workflows also opens new avenues for multimodal phenotyping. By combining cranial metrics with intracranial measurements – such as brain volume, cortical thickness, or ventricular size – clinicians and researchers can gain a comprehensive view of both skull and brain development^16^. This fusion of cranial and cerebral biomarkers enhances the ability to stratify disease risk, monitor progression, and evaluate treatment response in a range of pediatric neurological and neurodevelopmental disorders.

Here we present CranioTrace, a pipeline for automated quantification of head circumference and cranial shape from both research and routine clinical neuroimaging data. Of note, CranioTrace can be used for both MRI and CT data. In addition, using a large reference dataset, we provide population reference charts for the resulting phenotypes and calibrate these growth charts to specific neurogenetic syndromes. Both the imaging pipeline and the reference charts are available as a resource for other researchers and clinicians (link available upon publication). Together, these tools represent a significant advance in both the research and clinical domains for the potential of imaging-based, individualized assessments of head circumference and cranial shape.

## Results

We first evaluated whether CranioTrace could reliably extract cranial contours from brain MRI and CT scans. The pipeline consists of automated registration to age-appropriate templates, extraction of the external head contour across axial slices and three small head-pitch rotations, rejection of geometrically implausible contours, and retention of the largest passing circumference (Fig.1; see Methods). As described below in detail, the quality of automatically extracted cranial contours was demonstrated by high test-retest reliability, manual quality assessment, and agreement with tape measure-extracted measures of head circumference. We used population modeling to create usable references for imaging-derived head circumference, as well as other measures of cranial size, shape and asymmetry. Finally, the resulting models were applied to quantify deviations in groups with clinical cranial abnormalities.

**Figure 1:**
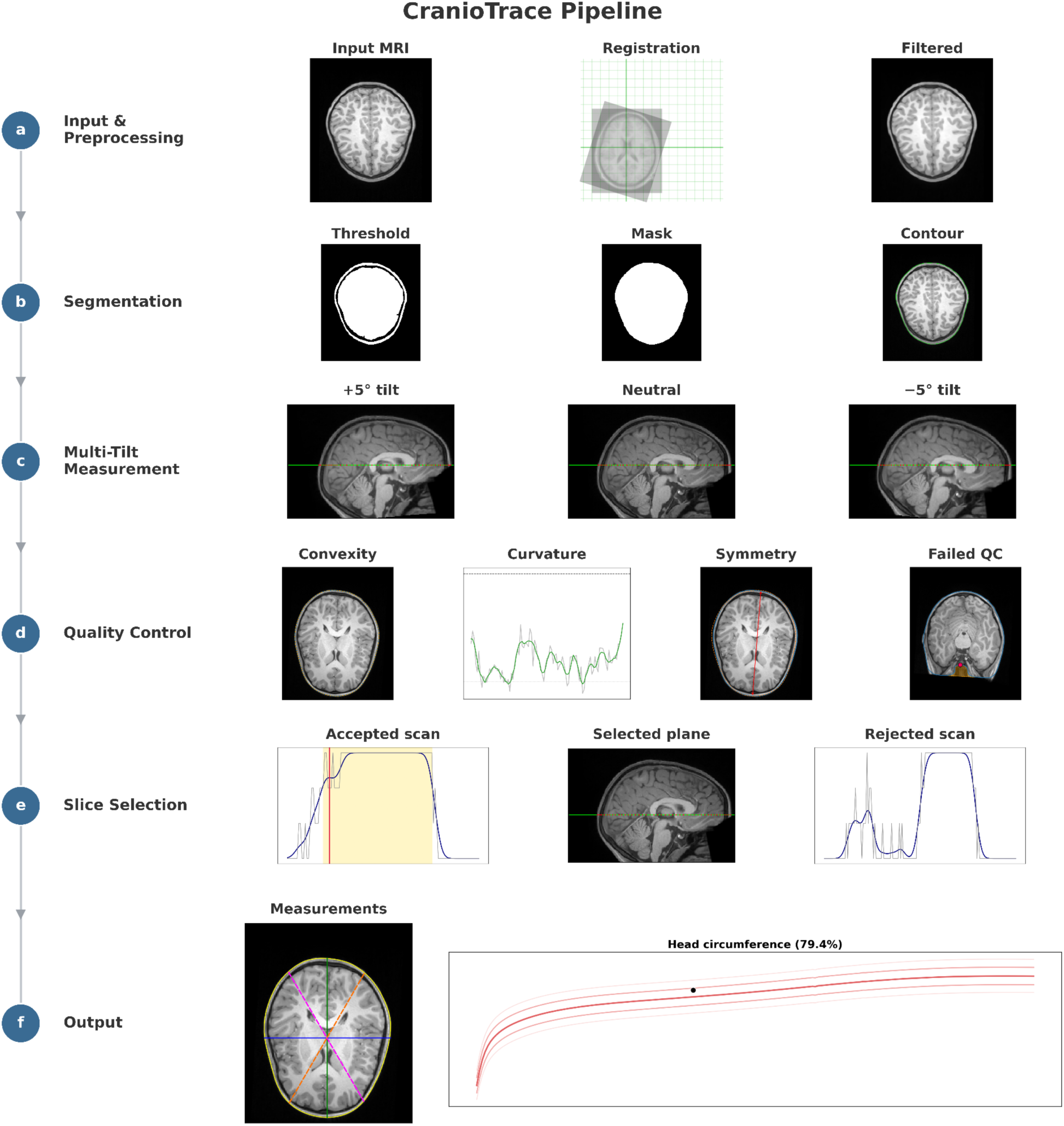
Overview of the CranioTrace pipeline for automated head-circumference and cranial-shape measurement from MRI. Each scan passes through six stages. **a,** Input and preprocessing: the T1- weighted scan is registered to an age-appropriate template and denoised with curvature anisotropic diffusion. **b,** Segmentation: the head is separated from the background by intensity thresholding and region growing, and an external cranial contour is extracted. **c,** Multi-tilt measurement: each volume is analyzed at three rotations (−5°, 0°, +5°) about the left–right axis to mitigate residual head tilt. **d,** Quality control: each contour is checked for convexity, curvature and bilateral symmetry, with failing scans flagged (a failed example is shown). **e,** Slice selection: an axial slice is chosen from the band of highest QC-pass ratio. **f,** Output: head circumference and cranial-shape indices, with population percentiles computed from the selected contour.

### Growth charts of head circumference

To establish reference charts for MRI-derived head circumference, we considered 20,556 structural MRI scans from 25 independent cohorts spanning birth to 41 years. Of these, 14,290 had the required metadata and met the eligibility criteria, 11,549 passed quality control, and 9,685 were included in the final analysis (Fig. S1). Of these, 8,968 were within the prespecified reporting range from birth to 20 years. We retained eligible observations beyond age 20 during model fitting to reduce upper-boundary effects and truncated the resulting reference curves at 20 (see Methods).

We then fitted sex-specific growth curves using generalized additive models for location, scale, and shape (GAMLSS) with a Box-Cox power exponential (BCPE) distribution – the distributional framework used to construct the WHO anthropometric head circumference-for-age standards (Fig. 2b)^1,17^. These models estimated age- and sex-dependent median trajectories and variability through nonlinear penalized splines. Subsequently, individual measurements can be benchmarked against the population model as centiles or centile z-scores.

**Figure 2:**
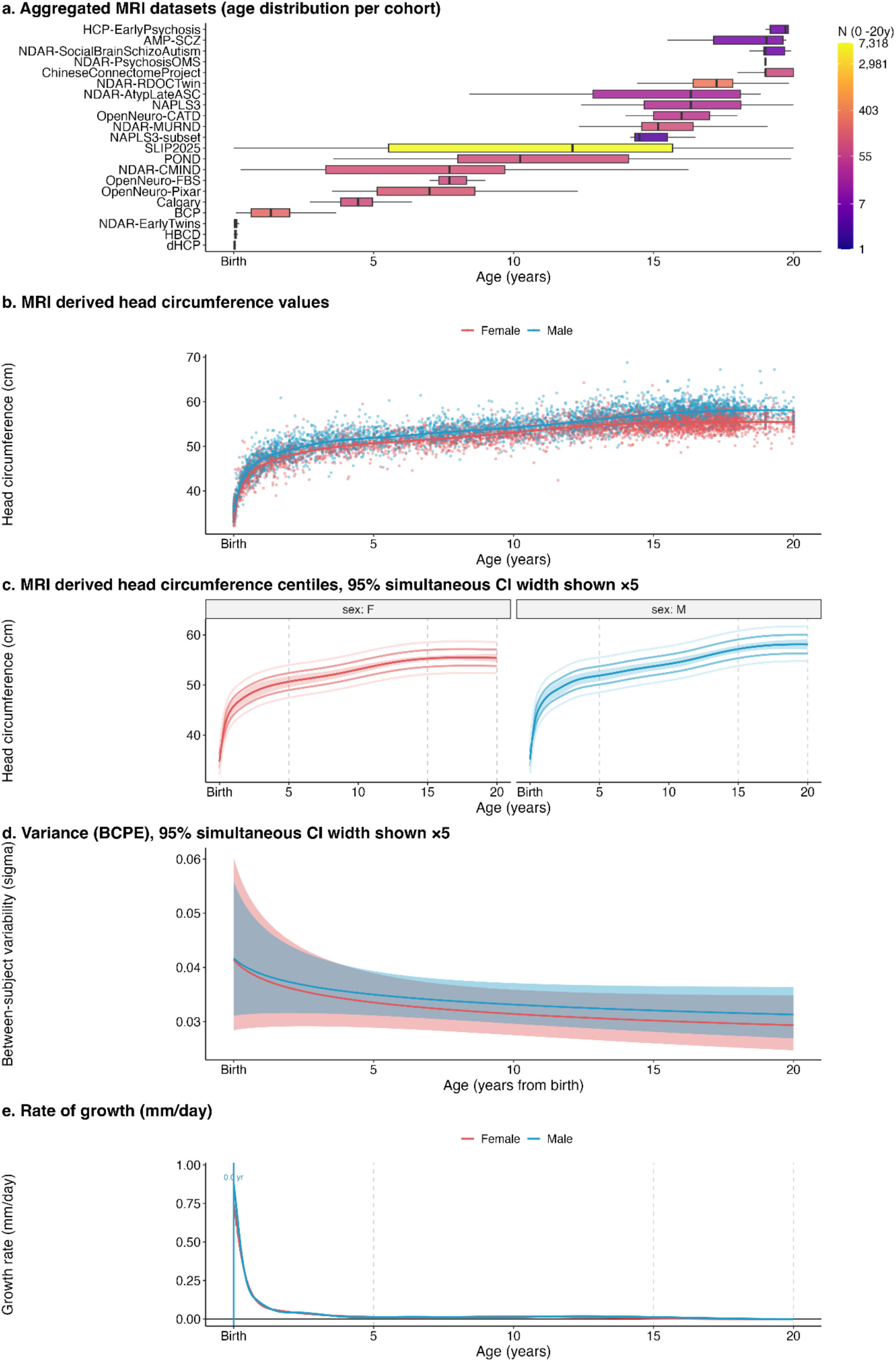
Age- and sex-specific trajectories of MRI-derived head circumference from birth to 20 years. **a,** Aggregated MRI data from 21 cohorts consisting of 8,968 scans with ages from birth to age 20. Box plots show the age distribution for each cohort where each box represents the interquartile range (IQR, 25th– 75th percentile), with a median central line and 95% whisker lines (2.5th–97.5th percentile). Box color illustrates sample size per cohort colored as the natural logarithm of n samples within the cohort. **b,** Head circumference measurements (cm) derived from the MRI pipeline along with median centile lines plotted against age stratified by sex (male, blue; female, red). **c,** GAMLSS estimated centile lines (3, 15, 50, 85, 97) for MRI derived head circumference measurements stratified by sex; 95% simultaneous CI bootstrapped with N = 200, magnified by a factor of 5 for visibility. **d,** Sex stratified median lines of estimated *σ* from the BCPE distribution which is approximately the coefficient of variation of the distribution with simultaneous CI from bootstrapping (N = 200). **e,** Rate of growth (mm/day), calculated as the difference in median head circumference between days.

As expected, MRI-derived head circumference followed a highly nonlinear trajectory, with rapid neonatal expansion giving way to asymptotic stabilization in adolescence (Fig. 2b). Males had larger median values than females, with non-overlapping 95% simultaneous confidence intervals after the immediate perinatal period (Fig. 2c). The BCPE scale parameter *σ*, which approximates relative between-subject variability, was highest at birth, decreased rapidly during the first five years, and plateaued by early adolescence (Fig. 2d). Growth velocity peaked at approximately 0.8 mm per day at birth, fell below 0.1 mm per day by age two, and approached zero by age five (Fig. 2e).

### Correlations with tape measurements and replicability of measurements

To assess agreement with the clinical standard for head circumferences, derived by tape measure, we compared MRI-derived head circumference (HC) with tape measurements in four cohorts where they were available for the same participants. Measurements were obtained in the same session for two cohorts (HBCD and dHCP, see Methods) and obtained within the same month for two cohorts (16p and SLIP2025, see Methods). Applying Bland-Altman analyses (Fig. 3), mean difference between MRI and tape measure in HBCD was 0.00 cm (s.d., 0.94, 95% limits of agreement (LoA), −1.84 to 1.85 cm); mean difference in 16p was −0.17 cm (s.d., 1.51; LoA, -3.13 to 2.78 cm); mean difference in SLIP2025 was 0.12 cm (s.d., 1.07; LoA, - 1.98 to 2.21 cm); and mean difference in dHCP was 0.13 cm (s.d., 0.75; LoA, −1.35 to 1.60 cm).

**Figure 3:**
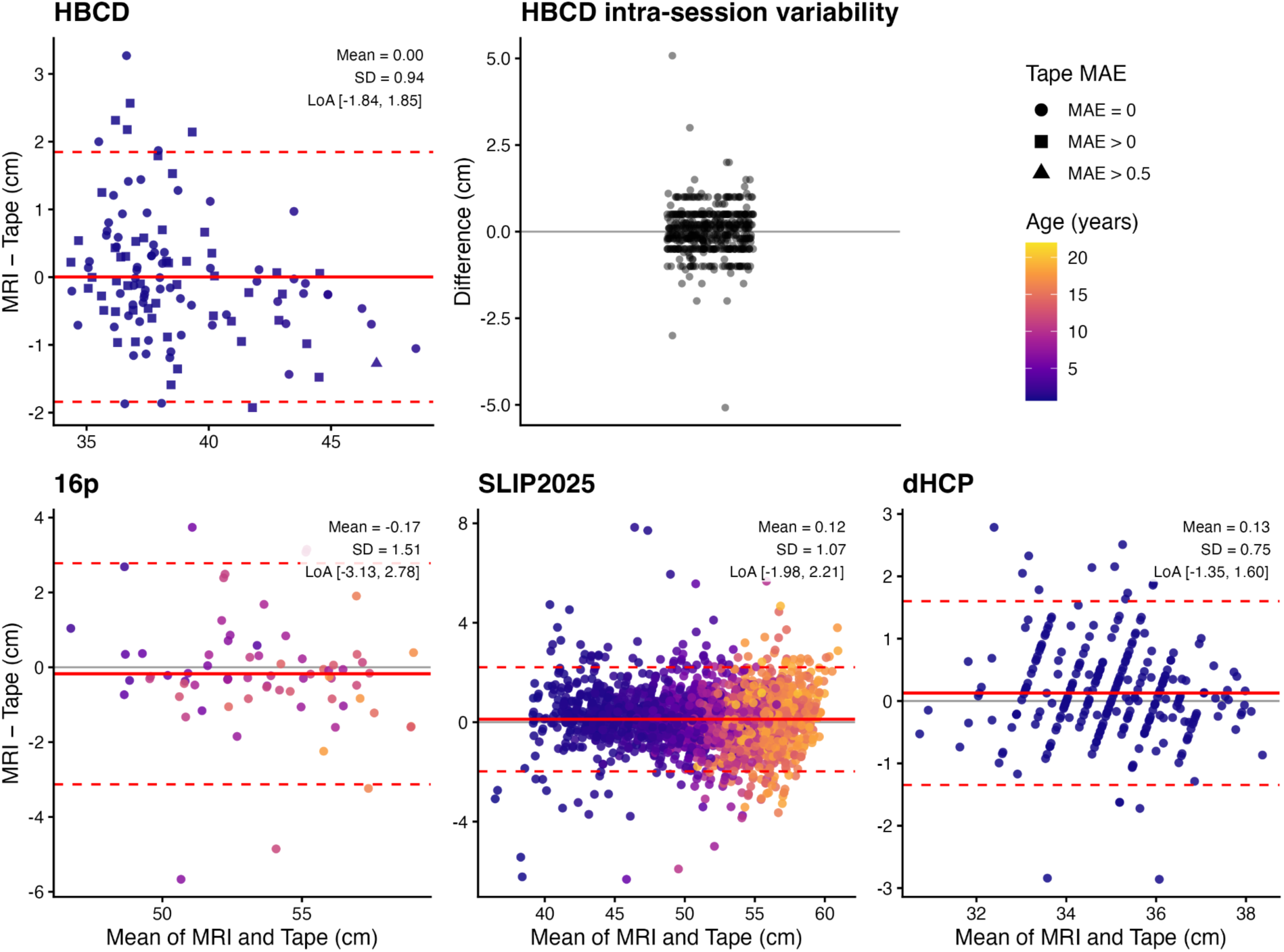
Agreement between head circumference estimates derived from magnetic resonance imaging (MRI) and physical tape measurements. Four datasets are visualized using Bland-Altman plots: the HEALthy Brain and Child Development study (HBCD; top left; n = 119 paired observations from 113 infants aged 5–229 days), a US multisite longitudinal study of early brain and child development; the 16p11.2 copy-number-variant cohort (bottom left; n = 71 paired observations from 67 youth aged 1.66–17.10 years with copy-number variants at the 16p11.2 locus, associated with microcephaly in duplication carriers and macrocephaly in deletion carriers); the SLIP2025 cohort (Scans with Limited Imaging Pathology, 2025 release; bottom middle; n = 2717 paired observations from 2535 Children’s Hospital of Philadelphia clinical-control participants aged from birth to 21.2 years with limited imaging pathology); and the Developing Human Connectome Project (dHCP; bottom right; n = 277 paired observations from 275 neonates aged 36.57–44.43 weeks postmenstrual age), an open neonatal neuroimaging study of perinatal brain development. The top-right panel illustrates HBCD intra-session variability in tape measurements; the largest difference among the available measurements in each included session is displayed symmetrically around zero. In each Bland-Altman plot, the y axis represents the difference between methods (MRI minus tape, cm), while the x axis represents the mean of the paired measurements. The solid red line indicates the mean bias, and the dashed red lines indicate the 95% limits of agreement. Data points are color-coded by age. Symbols in the HBCD cohort indicate the mean absolute error (MAE) among the intra-session tape measurements: circle, 0 cm; square, >0 to 0.5 cm; and triangle, >0.5 cm.

To assess CranioTrace’s test-retest reliability under realistic clinical conditions, we used a cohort of clinically acquired scans from the Children’s Hospital of Philadelphia (SLIP2025) with multiple high-resolution MPRAGE T1-weighted acquisitions within a single MRI session. Across 772 multi-scan sessions from 258 subjects, the unique scans were predominantly pre- and post- contrast (93.1%) and, less often, varied in acquisition parameters (voxel size, slice number). We converted each within-session MRI measurement to an age- and sex-referenced BCPE z-score before calculating intraclass correlations (ICCs) across repeat scans. Repeatability was high: ICC(1,1) = 0.98 and ICC(1,2) = 0.99. The within-session mean absolute error (MAE) was 0.07 z- score units, equivalent to 0.13 cm on the raw head-circumference scale. For comparison, we analyzed repeated same-day tape measurements which were acquired under optimized research conditions as part of the HBCD study. Of 636 study sessions, 622 had three measurements: two obtained by one examiner on the same occasion and a third obtained later that day by a second examiner. The remaining 14 sessions had two measurements from one examiner. After conversion with the CDC 2000 and UK-WHO growth references^1,18–20^, the within-session mean absolute error was 0.04 CDC z-score units, 0.05 UK-WHO z-score units, and 0.06 cm on the raw scale.

### Manual quality control assessment

To ensure the geometric integrity of the cranial outlines, we implemented an automated quality control (QC) pipeline evaluating shape convexity, curvature smoothness, and left–right symmetry (see Methods). Contours were required to pass strict quantitative thresholds across all three domains to successfully exclude artifacts like partial skull capture or residual extracranial tissue. The pipeline then selects the largest passing circumference within a single contiguous axial band. To manually assess the quality of slice selection and contour accuracy of the pipeline, one rater manually reviewed 1,000 scans from the CHOP SLIP cohort using the four-level rubric in Table 1. Overall, 973 scans (97.3%) passed. In all 25 scans rated 3 for contour failure, automated QC had excluded the defective slice and orientation, so the failure did not propagate to the final measurement. Most of the 27 scans rated 4 had registration errors that produced an anatomically incorrect orientation, including anterior–posterior commissure misalignment. These findings indicate that automated QC reliably excluded contour failures, although registration errors remained a source of slice-selection failure requiring manual review in a small subset of scans

**Table 1.** Distribution of Manual QC Ratings of 1000 scans for Contour and Slice Selection.

| Rating | Category | Pass? | N (%) |
| --- | --- | --- | --- |
| 1 | Passing contour for all orientations; selected slice for all three orientations | Yes | 683 (68.3%) |
| 2 | Passing contour for all orientations; optimal slice for at least one orientation | Yes | 265 (26.5%) |
| 3 | Contour failure for at least 1 orientation | Yes | 25 (2.5%) |
| 4 | Passing contour for all orientations; optimal slice not selected | No; likely registration errors not detected by auto-QC | 27 (2.7%) |
| 1 | Passing contour for all orientations; selected slice for all three orientations | Yes | 683 (68.3%) |
| 2 | Passing contour for all orientations; optimal slice for at least one orientation | Yes | 265 (26.5%) |
| Total |  |  | 1,000 (100%) |

### Correlations between MRI and CT

To assess cross-modality consistency, we applied the pipeline to paired CT and MRI scans available in the datasets CERMEP and SynthRAD (see Methods) and calculated CT-minus-MRI Bland-Altman differences for head circumference and the selected axial level (Fig. 4). In CERMEP (n = 24; age range 23–65), the mean head-circumference difference was −0.66 cm (s.d. = 0.29; 95% LoA, −1.23 to −0.09 cm; MAE, 0.67 cm), and the mean difference in axial level selected for circumference extraction was 0.60 cm (s.d. = 0.91; LoA, −1.19 to 2.39 cm; MAE, 0.89 cm). In SynthRAD (n = 128; age range 3–93), the corresponding differences were 0.42 cm for head circumference (s.d. = 0.84; LoA, −1.24 to 2.07 cm; MAE, 0.66 cm) and −0.31 cm for selected axial level (s.d. = 1.07; LoA, -2.42 to 1.8 cm; MAE, 0.84 cm).

**Figure 4:**
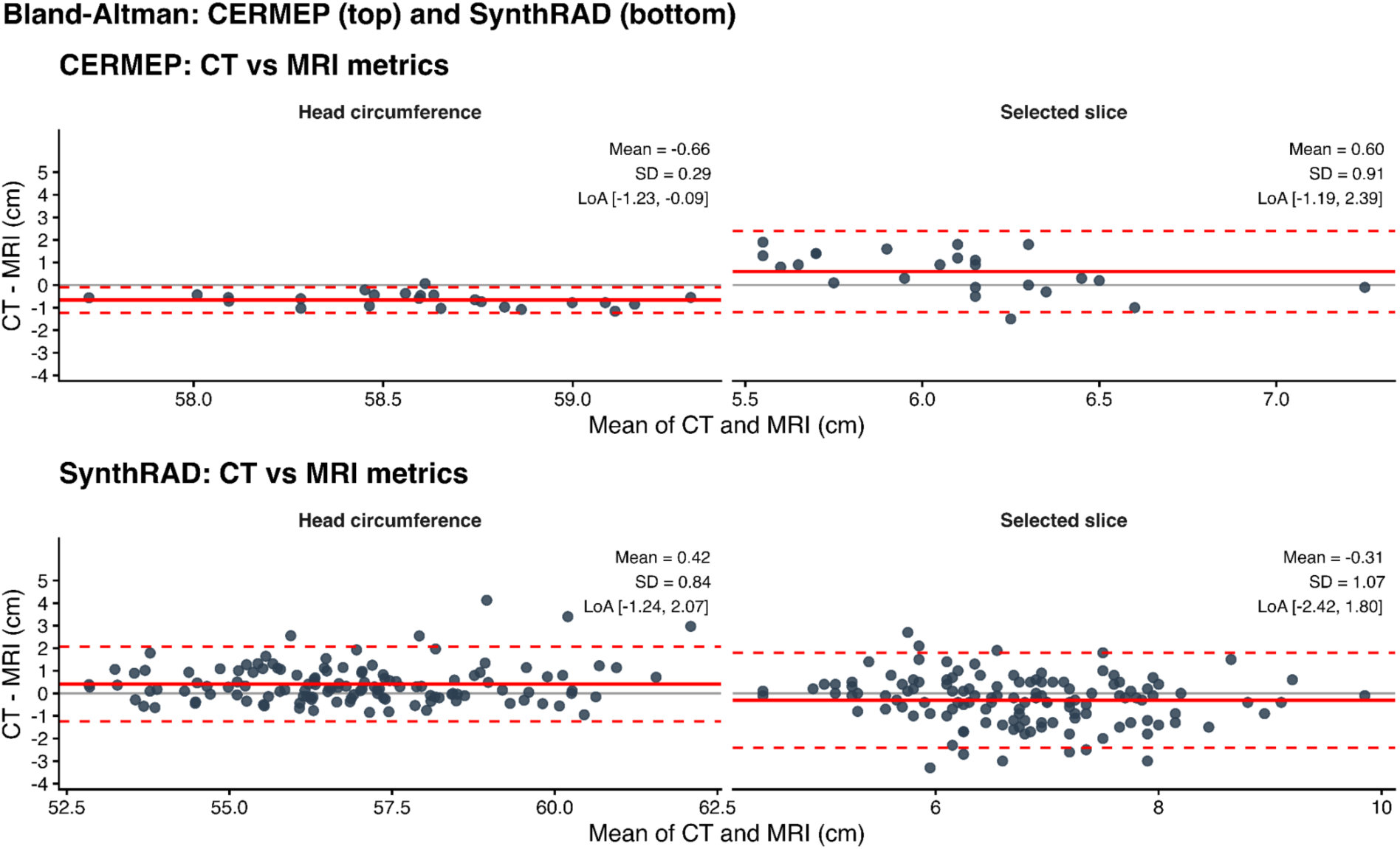
Bland-Altman agreement between CT- and MR-derived measurements in the CERMEP and SynthRAD cohorts. Bland–Altman plots of CT minus MR differences (cm) against the mean of CT and MR (cm) for head circumference (HC diff, left) and per-slice measurements (Slice diff, right) in the CERMEP cohort (top) and the SynthRAD cohort (bottom). Solid red lines indicate the mean difference; dashed red lines indicate the 95% limits of agreement (LoA). CERMEP: HC mean difference = −0.66 cm (s.d. = 0.29, LoA [−1.23, −0.09]); Slice mean difference = 0.60 cm (s.d. = 0.91, LoA [−1.19, 2.39]). SynthRAD: HC mean difference = 0.42 cm (s.d. = 0.84, LoA [−1.24, 2.07]); Slice mean difference = −0.31 cm (s.d. = 1.07, LoA [−2.42, 1.80]).

### Head circumference differences in neurogenetic cohorts

We tested the imaging-derived head-circumference measure in four neurogenetic conditions that affect head size. For each participant, the sex-specific BCPE reference model converted the measurement at the participant’s age to a z-score; zero denotes the reference median, and positive and negative values denote larger and smaller head circumference, respectively. We tested each cohort’s z-scores against zero using a one-sample Wilcoxon signed-rank test (Fig. 5).

**Figure 5:**
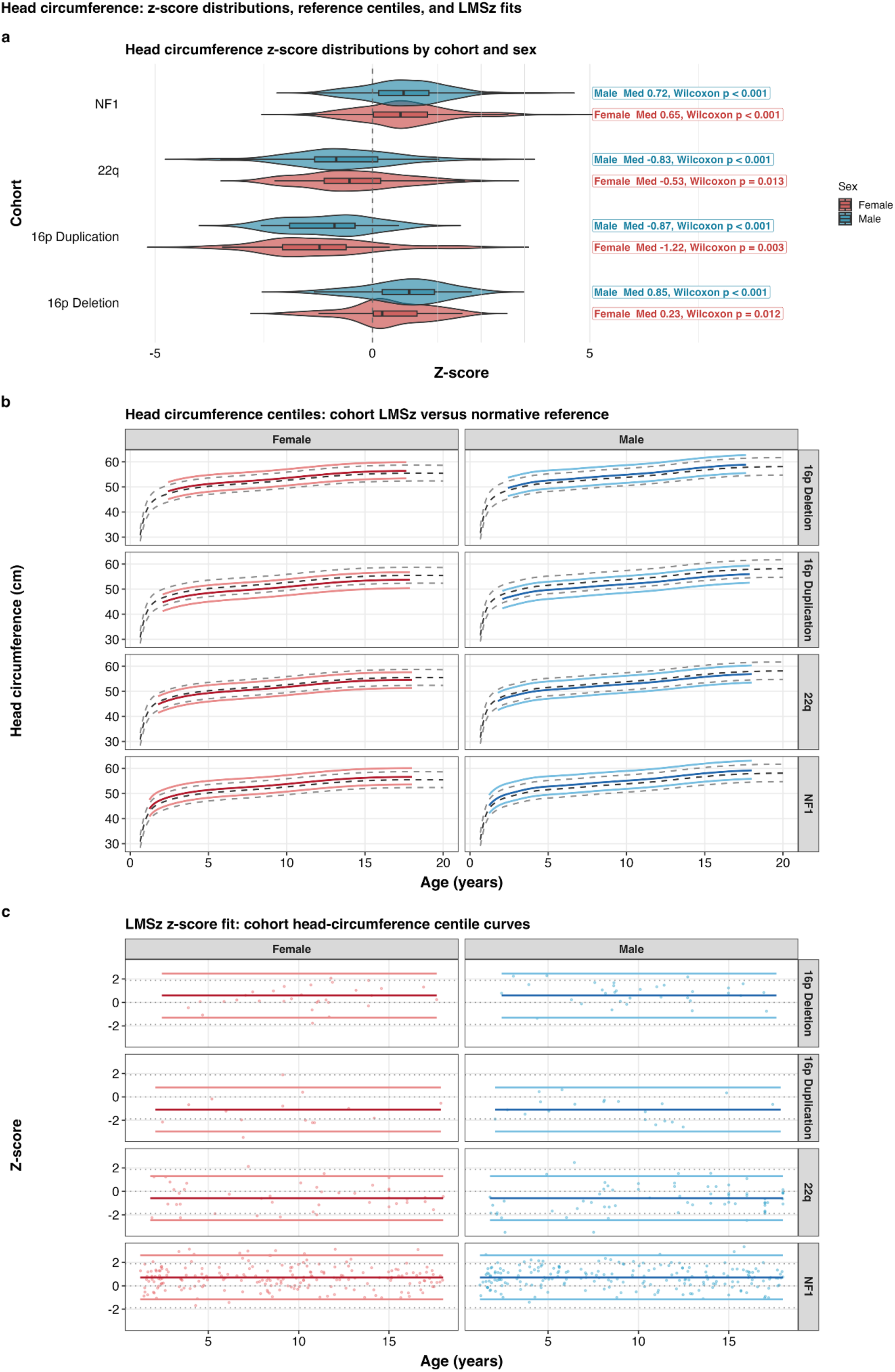
**a,** Split violin plots with embedded box plots show z-score distributions for the NF1, 22q11.2 deletion, 16p11.2 duplication, and 16p11.2 deletion cohorts, separated by sex (female, red; male, blue). Boxplots indicate the median and interquartile range; whiskers extend to 1.5× the interquartile range. The dashed vertical line at Z = 0 denotes the population reference mean. Positive shifts relative to zero are evident in NF1 and 16p deletion; negative shifts, in 22q and 16p duplication**. b,** Cohort-specific head circumference growth charts generated using the LMSz method^21^. Curves are back-transformed centile lines (3rd, 50th, and 97th) for females (red) and males (blue) from early childhood through late adolescence, plotted alongside the corresponding centiles for the non-clinical reference cohort (grey). **c,** Z-score fits with cohort centile lines at z = −2, 0, and +2, with individual observations overlaid.

Despite substantial within-condition heterogeneity, head circumference was increased in the macrocephaly-associated conditions. In both NF1 and 16p11.2 deletion syndrome, HC was elevated in both females (NF1: n = 207, median z = 0.65, IQR 0.02 to 1.27, range −1.69 to 4.29, p < 0.001; 16p11.2: n = 30; 0.23, IQR 0.02 to 1.03, range −1.77 to 2.07; p = 0.012) and males (NF1: n = 207, 0.72, IQR 0.15 to 1.30, range −1.40 to 3.84, p < 0.001; 16p11.2: n = 33; 0.85, IQR 0.23 to 1.43, range −1.35 to 2.29; p < 0.001).

In contrast, head circumference was decreased in conditions commonly associated with microcephaly. In both 16p11.2 duplication and 22q11.2 deletion syndromes, HC was reduced in females (16p11.2: n = 15; −1.22, IQR −2.07 to −0.61, range −3.46 to 1.88; p = 0.003; 22q: n = 43; −0.53, IQR −1.11 to 0.19, range −2.25 to 2.13; p = 0.013) and males (16p: n = 20; −0.87, IQR −1.90 to −0.40, range −2.57 to 0.61; p < 0.001; 22q: n = 65; −0.83, IQR −1.33 to 0.12, range −3.50 to 2.46; p < 0.001).

We next used the LMSz method to convert each condition’s distribution of reference z-scores into condition-specific growth charts^21^ (Fig. 5b, c). The microcephaly-associated cohorts (16p11.2 duplication and 22q11.2 deletion) showed negative offsets from the reference curves, whereas the macrocephaly-associated cohorts (NF1 and 16p11.2 deletion) showed positive offsets. In all four cohorts, the best-fitting secondary model used constant location and scale parameters; the estimated offset therefore did not vary by age or sex.

### Measures of Cranial Shape and Asymmetry

#### Age-related changes in cranial shape and asymmetry

To characterize cranial morphology beyond global size, we derived eight head-shape metrics from the automatically selected axial contour and fitted sex-specific, age-dependent GAMLSS reference curves (Fig. 6; Table 2). The metrics captured cranial proportion and roundness (cephalic index (CI), anterior–posterior ratio (APR), and cranial roundness index (CRI)), diagonal vault asymmetry (cranial vault asymmetry index (CVAI)), and directional left–right asymmetry (asymmetry coefficient (AC), anterior hemisphere asymmetry (AHA), posterior hemisphere asymmetry (PHA), and left hemisphere ratio (LHR)). We retained the signs of the directional metrics rather than collapsing them to absolute magnitudes. Positive AC, AHA, and PHA values therefore indicate a larger left-sided area, whereas negative values indicate a larger right-sided area; LHR > 1 likewise indicates a larger left-sided area.

**Figure 6:**
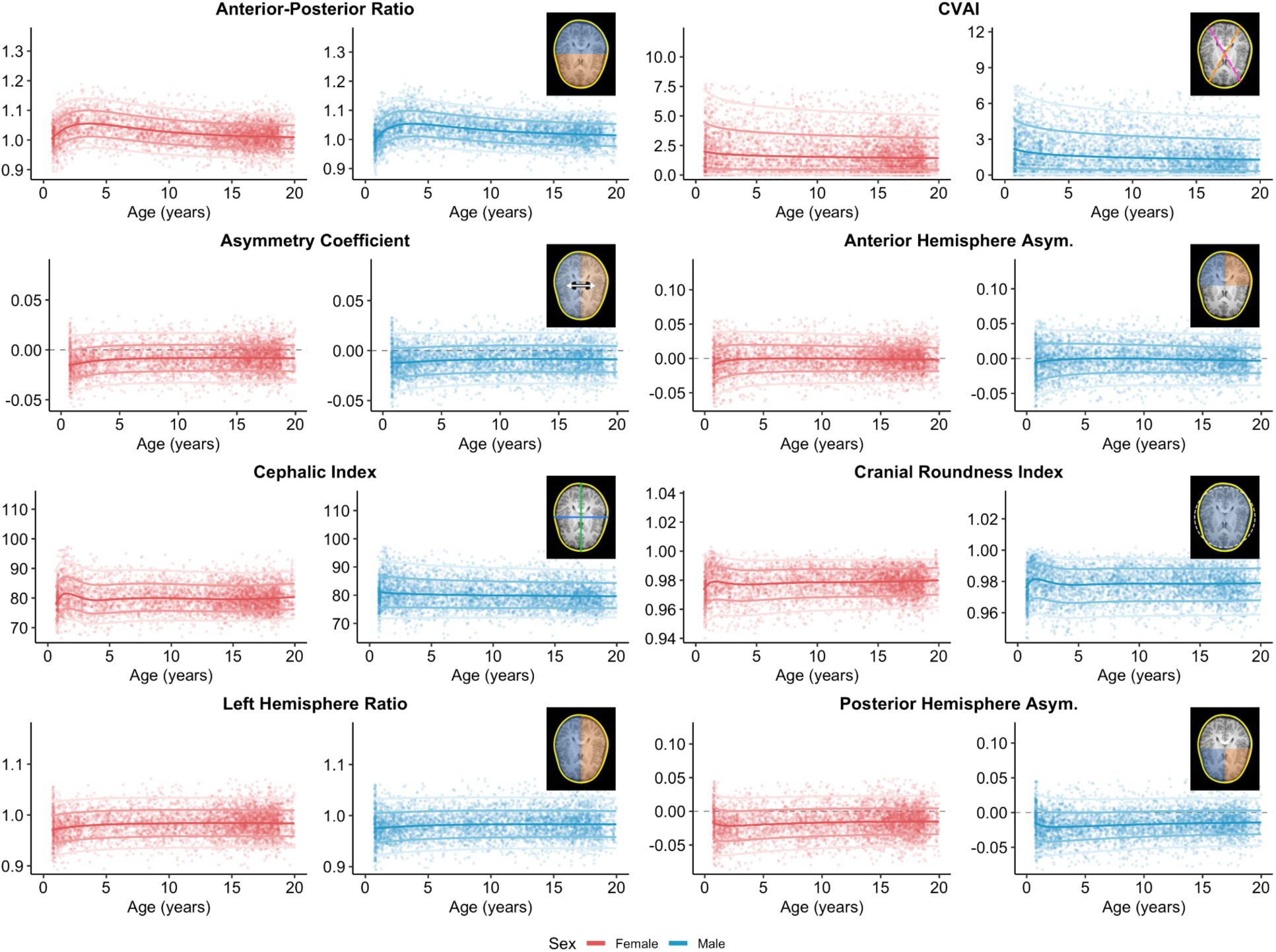
Reference centile curves for cranial shape and asymmetry metrics. The panels display growth trajectories from early childhood through late adolescence for females (red) and males (blue) across eight morphological indices: Cephalic Index, Cranial Vault Asymmetry Index, Asymmetry Coefficient, Anterior Hemisphere Asymmetry, Anterior–Posterior Ratio, Cranial Roundness Index, Left Hemisphere Ratio, and Posterior Hemisphere Asymmetry. The solid curves represent the 3rd, 15th, 50th (bold), 85th, and 97th centiles, with individual scan observations overlaid.

**Table 2.** Measures of cranial shape and asymmetry. For each metric we report its equation, the geometric quantity it measures, and the cranial feature or condition it indexes. Equations are reproduced from the Methods for reference. Quadrant areas are defined relative to the geometric center of the contour’s bounding box: AL, anterior-left; AR, anterior-right; PL, posterior-left; PR, posterior-right; total area is the full area enclosed by the contour. In CI, Width and Length are the bounding-box dimensions; in CVAI, the longer and shorter diagonals are the diameters measured at 60° and 120° through the centroid. Linear dimensions are in millimeters (mm) and areas in square centimeters (cm^2^); CI and CVAI are expressed as percentages, and AC, AHA, PHA, CRI, LHR, and APR are dimensionless. For the signed metrics, positive values denote a left-sided excess (AC, AHA, PHA); likewise, LHR > 1 indicates left-dominant area and APR > 1 indicates anterior-dominant area.

| Metric | Equation | What it measures | What it indexes |
| --- | --- | --- | --- |
| Cephalic Index (CI) | $(\text{Width} / \text{Length}) \times 100$ | How wide the head is relative to its length, as a percentage. | Overall head proportion; high values indicate brachycephaly (short/wide), low values indicate dolichocephaly (long/narrow). |
| Cranial Vault Asymmetry Index (CVAI) | $((\text{Longer Diagonal} - \text{Shorter Diagonal}) / \text{Longer Diagonal}) \times 100$ | How much the two diagonal diameters differ. | Oblique (diagonal) skull distortion; the standard index of plagiocephaly. |
| Asymmetry Coefficient (AC) | $((\text{AL} + \text{PL}) - (\text{AR} + \text{PR})) / \text{total\_area}$ | Whether the left and right halves enclose equal area; sign shows which side is larger. | Global left–right area imbalance (directional). |
| Anterior Hemisphere Asymmetry (AHA) | $(\text{AL} - \text{AR}) / (\text{AL} + \text{AR})$ | Within the front of the head, how unequal the left and right sides are. | Regional (frontal) left–right asymmetry; anterior/forehead plagiocephaly. |
| Posterior Hemisphere Asymmetry (PHA) | $(\text{PL} - \text{PR}) / (\text{PL} + \text{PR})$ | Within the back of the head, how unequal the left and right sides are. | Regional (occipital) left–right asymmetry; posterior deformational plagiocephaly. |
| Cranial Roundness Index (CRI) | $(4 \times \pi \times \text{total\_area}) / \text{circumference\_mm}^2$ | How close the outline is to a perfect circle (1.0 = circle). | Overall roundness; circularity of the cranial outline. |
| Left Hemisphere Ratio (LHR) | $(\text{AL} + \text{PL}) / (\text{AR} + \text{PR})$ | Ratio of left-side area to right-side area (1.0 = balanced). | Hemispheric area balance; >1 indicates leftward dominance. |
| Anterior–Posterior Ratio (APR) | $(\text{AL} + \text{AR}) / (\text{PL} + \text{PR})$ | Ratio of front area to back area (1.0 = balanced). | Anteroposterior balance; frontal–occipital taper, larger means more frontally biased. |

Across the reference sample age window (n = 9,685), most cranial shape metrics showed their strongest age-related changes in infancy and early childhood (Fig. 6). CVAI was highest early in life and decreased with age, consistent with attenuation of oblique cranial vault asymmetry over development. CI and APR showed early-life peaks followed by gradual stabilization or decline, indicating changes in width-length and anterior–posterior proportions as the skull expands. CRI increased rapidly in infancy, showed a modest early dip, and then increased gradually, reflecting developmental changes in overall cranial roundness.

The signed asymmetry metrics showed that directional information was not redundant with unsigned asymmetry magnitude. In the fitted reference curves, global left–right balance remained close to symmetric, and signed AHA stayed centered near zero across development. By contrast, the model-estimated median trajectory for signed PHA was consistently shifted below zero across much of the age range, indicating a right-posterior predominance in the reference model. This signed formulation allows anterior and posterior asymmetry to be interpreted as regional directional components of cranial torque in individuals, although the fitted population-level pattern was driven more by posterior rightward asymmetry than by a strong global left-anterior/right-posterior opposition.

Together, these results show that cranial shape and directional asymmetry follow structured developmental trajectories that are not captured by head circumference alone. The most rapid changes occur during the same early-life period in which head circumference grows fastest, but several metrics retain stable age- and sex-specific reference distributions into later childhood and adolescence.

#### Asymmetry differences between Cranial Shape Groups

We next tested whether conventional cephalic-index categories were associated with distinct age- and sex-adjusted cranial morphology (Fig. 7; Table 3). Raw CI classified scans with finite z-scores for all modelled metrics as dolichocephalic (CI < 75; n = 1,096), mesocephalic (75 <= CI < 80; n = 3,779), or brachycephalic (CI >= 80; n = 4,810). We compared z-scores between groups using linear mixed-effects models with study as a random intercept and mesocephalic scans as the reference; models of non-HC metrics also adjusted for HC z-score. Although CI varies nonlinearly with age, its use here was limited to conventional descriptive grouping, whereas all outcome comparisons used age- and sex-referenced z-scores.

**Figure 7.**
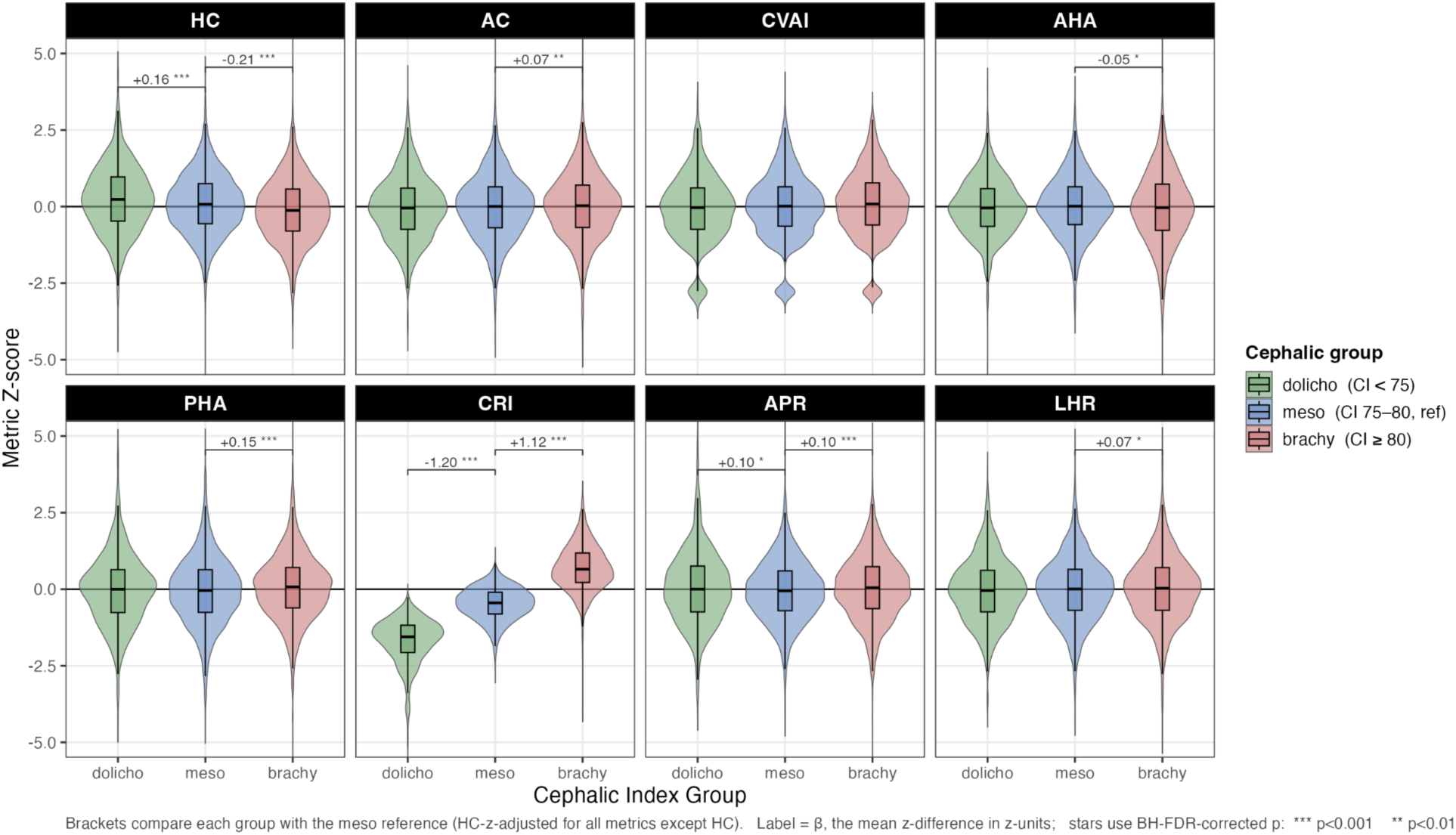
Cranial features across cephalic index groups. Cranial morphology z-scores across cephalic index groups. Violin and box plots show age- and sex-referenced z-scores for head circumference (HC), asymmetry coefficient (AC), cranial vault asymmetry index (CVAI), anterior hemisphere asymmetry (AHA), posterior hemisphere asymmetry (PHA), cranial roundness index (CRI), anterior–posterior ratio (APR), and left hemisphere ratio (LHR), stratified by conventional raw cephalic index categories: dolicho (CI < 75), meso (75 <= CI < 80, reference), and brachy (CI >= 80). Brackets compare each group with the mesocephalic reference group; labels report the linear mixed-effects beta coefficient in z-units and BH-FDR corrected significance level. Non-HC models are adjusted for HC-z and include a random intercept for study.

**Table 3:** Linear mixed-effects models testing associations between cephalic index group and cranial morphology z-scores. Mesocephalic scans are the reference group. For non-HC metrics, beta coefficients for dolicho and brachy groups are adjusted for HC-z and study random intercepts. The HC-z column reports the covariate association from the same additive model. All reported p-values are Benjamini- Hochberg FDR-corrected across a single family of the 37 model terms in this table.

| Outcome z-score | Dolicho vs meso (Est.) | Dolicho vs meso (p) | Brachy vs meso (Est.) | Brachy vs meso (p) | HC-z assoc. (Est.) | HC-z assoc. (p) | Dolicho x HC-z (Est.) | Dolicho x HC-z (p) | Brachy x HC-z (Est.) | Brachy x HC-z (p) |
| --- | --- | --- | --- | --- | --- | --- | --- | --- | --- | --- |
| HC | +0.16 | < 0.001 | -0.21 | < 0.001 | N/A | N/A | Not tested | Not tested | Not tested | Not tested |
| AC | -0.03 | 0.509 | +0.07 | 0.005 | +0.03 | 0.015 | +0.01 | 0.776 | -0.04 | 0.136 |
| CVAI | -0.06 | 0.143 | +0.02 | 0.412 | -0.13 | < 0.001 | +0.03 | 0.412 | +0.03 | 0.303 |
| AHA | -0.06 | 0.158 | -0.05 | 0.049 | +0.02 | 0.158 | +0.05 | 0.244 | +0.01 | 0.789 |
| PHA | +0.02 | 0.758 | +0.15 | < 0.001 | +0.02 | 0.059 | -0.03 | 0.451 | -0.06 | 0.018 |
| CRI | -1.20 | < 0.001 | +1.12 | < 0.001 | -0.05 | < 0.001 | -0.07 | 0.005 | -0.02 | 0.179 |
| APR | +0.10 | 0.017 | +0.10 | < 0.001 | -0.07 | < 0.001 | +0.09 | 0.015 | -0.00 | 0.975 |
| LHR | -0.03 | 0.465 | +0.07 | 0.010 | +0.03 | 0.016 | +0.01 | 0.776 | -0.04 | 0.158 |

Head circumference deviation was associated with several cranial morphology metrics after accounting for CI group and study (Table 3). Higher HC-z was associated with lower CVAI-z, CRI-z, and APR-z, and with modestly higher AC-z, and LHR-z. Neither AHA-z nor PHA-z was significantly associated with HC-z. Secondary interaction models showed that the HC-z association differed by CI group only for brachycephalic PHA-z, dolichocephalic CRI-z, and dolichocephalic APR-z.

CI-group effects were dominated by roundness and cranial proportion. Compared with mesocephalic scans, dolichocephalic scans had markedly lower CRI-z, whereas brachycephalic scans had markedly higher CRI-z. Both dolichocephalic and brachycephalic scans also showed modestly higher APR-z, while CVAI-z did not differ significantly between CI groups after HC-z adjustment.

Signed asymmetry effects were smaller and were concentrated in the brachycephalic group. Compared with mesocephalic scans, brachycephalic scans showed shifts in signed AC-z, AHA- z, PHA-z, and LHR-z, whereas dolichocephalic scans did not differ significantly from mesocephalic scans for any signed asymmetry metric.

Overall, conventional CI groups primarily separated along roundness and anterior–posterior proportionality, with smaller accompanying shifts in signed asymmetry. These findings support treating head circumference, cephalic proportion, roundness, anterior–posterior balance, and directional asymmetry as related but nonredundant components of cranial morphology.

#### Cranial shape differences in neurogenetic cohorts

To determine whether the neurogenetic cohorts showed cranial-shape differences beyond those expected from head circumference, we compared eight head-circumference-conditioned shape z scores with the reference expectation of zero (Fig. 8; Supplementary Table 1). Only NF1 showed shape deviations that survived Benjamini–Hochberg correction. Among participants with NF1 (n = 414), the mean conditional z scores were −0.12 for cephalic index (95% confidence interval, −0.21 to −0.03; adjusted P = 0.044) and 0.19 for cranial vault asymmetry index (0.09–0.28; adjusted P = 9.6 × 10^-4^). NF1 was also associated with lower cranial roundness (mean conditional z, −0.18; 95% confidence interval, −0.27 to −0.09; adjusted P = 8.2 × 10^-4^) and greater anterior weighting (mean conditional anterior–posterior ratio z, 0.35; 95% confidence interval, 0.25–0.46; adjusted P = 4.6 × 10^-10^). No additional shape comparison survived correction (all adjusted P ≥ 0.077).

**Figure 8.**
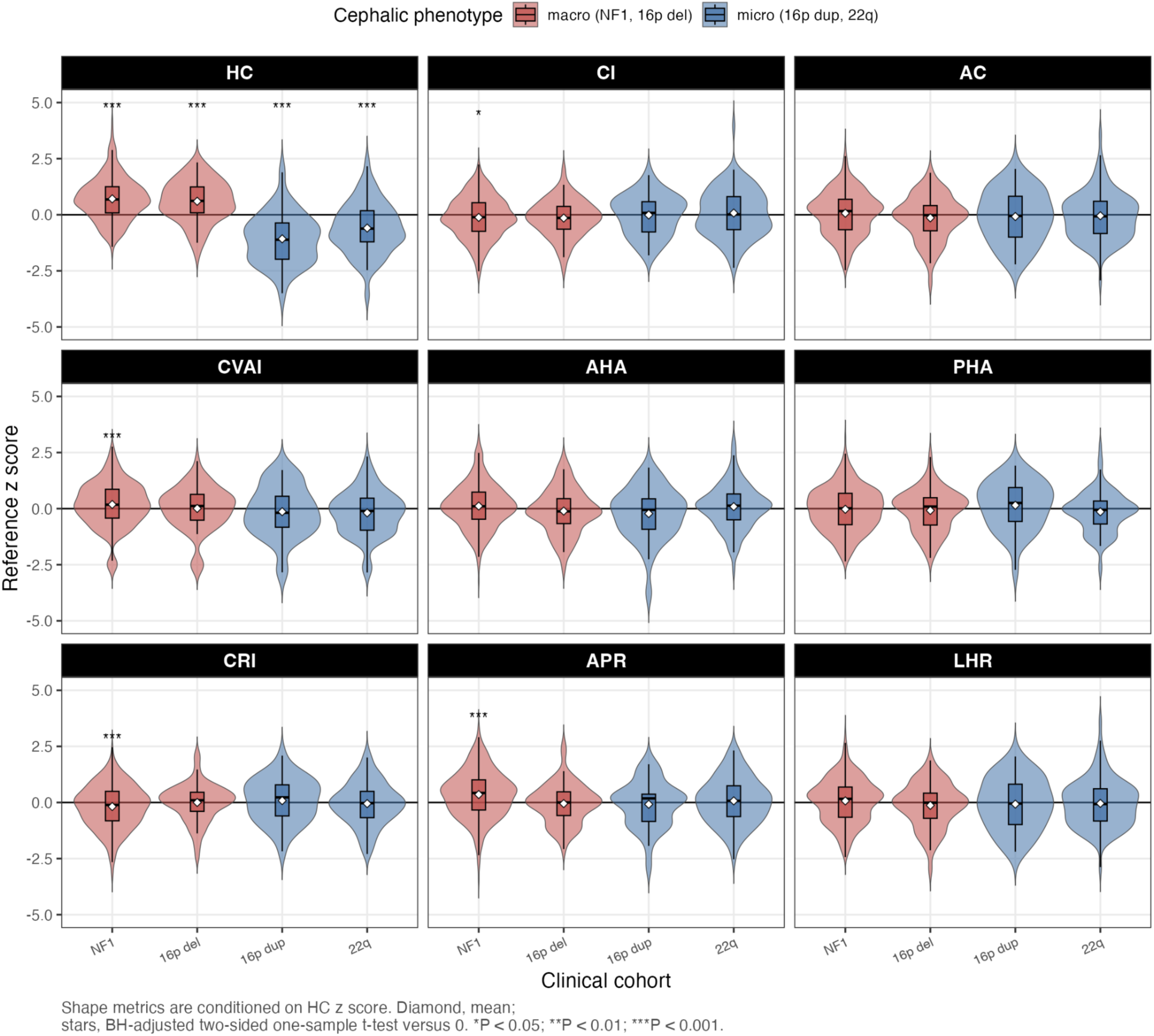
Head-circumference and cranial-shape deviations in four neurogenetic cohorts. Violin and embedded box plots show reference z-score distributions for NF1 (n = 414), 16p11.2 deletion syndrome (n = 63), 16p11.2 duplication syndrome (n = 35), and 22q11.2 deletion syndrome (n = 108). The head-circumference (HC) panel shows age- and sex-referenced z scores. The remaining panels show shape z scores conditioned on the linear association with HC z score in the reference sample and standardized by the corresponding reference residual standard deviation; zero therefore denotes the shape expected for an individual’s age, sex, and HC deviation. Boxes show the median and interquartile range, whiskers extend to 1.5 times the interquartile range, and white diamonds denote means. Red indicates conditions conventionally associated with macrocephaly (NF1 and 16p11.2 deletion), and blue indicates conditions conventionally associated with microcephaly (16p11.2 duplication and 22q11.2 deletion). Asterisks denote Benjamini–Hochberg-adjusted P values from two-sided one-sample t tests against zero, corrected across all 36 cohort–metric comparisons. One, two, and three asterisks denote adjusted P < 0.05, adjusted P < 0.01, and adjusted P < 0.001, respectively. AC, asymmetry coefficient; AHA, anterior hemisphere asymmetry; APR, anterior–posterior ratio; CI, cephalic index; CRI, cranial roundness index; CVAI, cranial vault asymmetry index; LHR, left hemisphere ratio; PHA, posterior hemisphere asymmetry.

## Discussion

We present a neuroimaging-based framework for quantifying head circumference, cranial shape, and cranial asymmetry across development, modelling age- and sex-specific reference curves from birth through 20 years of age. Using structural MRI from 9,685 individuals across 25 studies, we derived population centiles that recapitulated the expected pattern of rapid postnatal cranial expansion followed by marked deceleration through childhood and stabilization in adolescence. The resulting charts extend the logic of conventional tape-measured head circumference growth standards into the imaging domain, incorporating measures of shape and asymmetry that tape-based assessment cannot provide^1,2,12,13^.

Several aspects of the reference models have important biological implications. The scale parameter of the distribution (σ) – approximately the coefficient of variation^22^ – peaked at birth and declined through early childhood before plateauing in adolescence. In other words, the population becomes more homogeneous relative to its own median, with the distribution of percentage deviations tightening as children age. Growth velocity peaked at approximately 0.8 mm per day at birth and fell below 0.1 mm per day by age 2, tapering toward zero by age 5.

Together, these observations characterize the neonatal and infant periods as a window of both high inter-individual heterogeneity and rapid growth, consistent with the dynamic brain and skull maturation described in prior work^12,15^.

Our analyses suggest that MRI-derived head circumference can serve as a practical complement to traditional tape measurement. Across four cohorts, mean bias relative to tape was small (−0.17 to 0.13 cm). Moreover, intrasession MRI reproducibility in the CHOP SLIP cohort was high, with intraclass correlation coefficients of 0.98–0.99 and a mean absolute error of 0.13 cm in the repeated-scan analysis. Manual review of 1,000 outputs showed a 97.3% overall success rate. And importantly, most failures reflected registration problems rather than major contouring errors (which are addressed through the automatic QC), which could be addressed through further optimization of the preprocessing pipeline.

Comparing repeated MRI or tape-based head circumference measured on the same day, per-measurement precision was comparable between the two methods. Tape mean absolute error (MAE) in HBCD was 0.06 cm, marginally smaller than the same-day MRI MAE of 0.13 cm.

However, repeated tape measurements in HBCD were highly protocolized, unlike typical clinical tape measurements, which drift with operator technique, hair, patient cooperation, and craniofacial anatomy. The biggest value of the present approach may lie in standardized, reproducible measurement extracted from scans already being acquired – particularly when retrospective comparison scans are also available or when physical examination data are missing, noisy, or difficult to obtain.

Adaptability of the models across MRI and CT support the translational potential of the framework. This cross-modal applicability matters for clinical deployment because MRI and CT are used in different pediatric settings and age groups, and a modality-flexible approach raises the likelihood that cranial metrics can be extracted from existing archives. Nevertheless, it is important to note that MRI- and CT-derived values do require harmonization. In CERMEP, CT-derived head circumference was smaller than MRI-derived head circumference on average (mean difference −0.66 cm; LoA −1.23 to −0.09 cm), whereas in SynthRAD the bias ran in the opposite direction (mean difference 0.42 cm; LoA −1.24 to 2.07 cm). Cross-modal slice selection was likewise dataset-dependent, with a positive mean slice difference in CERMEP (0.60 cm; LoA −1.19 to 2.39 cm) and a slight negative mean difference in SynthRAD (-0.31 cm; LoA -2.42 to 1.80 cm). The cohort-specific direction of bias indicates that cross-modal differences depend on acquisition characteristics and image contrast rather than reflecting a single universal offset. A visual inspection of the samples in SynthRAD with a high positive difference revealed tissue-related differences between the two modalities. Direct comparison of a patient’s MRI- and CT-derived measurements should be interpreted cautiously, with modality-specific calibration required before MRI- and CT-derived values are combined in a single reference model.

Beyond the nearly ten thousand participants included in our MRI reference model, our results also illustrate how imaging-derived head circumference can support precision phenotyping in specific neurogenetic cohorts. Average deviations in neurogenetic cohorts followed the expected directions of effect, with negative shifts for cohorts with microcephalic tendencies (16p duplication and 22q deletion) and positive shifts for cohorts with macrocephalic tendencies (NF1 and 16p deletion). Notably, we also generated cohort-specific reference models even for relatively small-n neurogenetic cohorts, by borrowing strength from the reference distribution via the LMSz framework^21^. The availability of cohort-specific reference models may prove valuable when studying conditions in which abnormal cranial growth is a core but age-dependent component of the phenotype, because it incorporates disorder-specific deviations not only in absolute size but also in the timing, dispersion, and persistence of those deviations across development. Moreover, how an individual compares to a genetically informed reference may have more clinical relevance than comparing an individual with a neurogenetic syndrome to a reference derived from the healthy population^23,24^.

An additional contribution of this work is the demonstration that cranial shape and asymmetry can be charted alongside head circumference. Most asymmetry metrics changed most rapidly in infancy and early childhood before stabilizing, suggesting that population cranial remodeling is concentrated in the same developmental window as rapid head growth. These trajectories are plausibly relevant to disorders in which skull shape carries diagnostic or mechanistic information^10,11^. As shown by our analysis of the cephalic index, head size and head shape provide related but not redundant information. Even after normalization, head-circumference deviation independently moderated several asymmetry features, indicating that cranial morphology cannot be fully summarized by circumference alone. This argues for a multidimensional cranial phenotype in which overall size, roundness, anterior–posterior balance, and hemispheric asymmetry each contribute complementary information.

In the neurogenetic analyses, this multidimensional representation revealed an NF1 pattern beyond the linear size–shape relation observed in the reference sample. Lower cephalic index and cranial roundness, together with higher cranial vault asymmetry and anterior–posterior ratio, indicated a lower width-to-length ratio, less circular outline, greater diagonal asymmetry, and greater anterior weighting. No other cohort showed shape deviations that survived false-discovery-rate correction. Because the NF1 sample was substantially larger, however, the null findings should not be interpreted as evidence that cranial shape is unaffected in the smaller cohorts.

There are some methodological limitations within the present study that should be noted. First, a large portion of our reference data comes from clinically acquired scans from clinical controls, who are ascertained differently than scans of “healthy controls” in research studies. However, we have previously demonstrated high overlap in brain imaging-derived measurements between clinical controls and research controls^13^. Furthermore, the pipeline uses 2d contour tracing, relying on accurate slice selection after registration to a template, as opposed to 3d models which could in theory circumnavigate these requirements. However, we argue that the relative simplicity of the model is a major strength, allowing for easy troubleshooting of quality control issues. Finally, we present reference growth curves for neurogenetic syndromes, though the sample sizes are small for population-representative population models. However, the LMSz method is adapted to this challenge by borrowing strength from the control reference models rather than constructing new models from scratch for each neurogenetic syndrome^21^.

Despite these limitations, the present effort represents a significant step towards usable, neuroimaging-based measurements of head circumference and other craniofacial features. Critically, new measures are paired with population-scale models, allowing benchmarking of individual patients to reference charts. Both the image processing pipelines and the reference models are publicly available for non-commercial use by other researchers at https://github.com/BGDlab (upon publication).

## Methods

The overall pipeline for automated head circumference, area, and volume measurement from MRI scans is initiated by pre-processing the input MRI, which includes age-specific template registration and image normalization. Throughout, age denotes post-menstrual age corrected for prematurity wherever gestational age was available. Subsequently an iterative process segments the head contour on sequential slices, from which various morphometric measurements are derived. All code will be available upon publication, as well as containerized software, for ease of implementation.

### Children’s Hospital of Philadelphia dataset

As previously described^13,25–27^, clinical controls were aggregated from brain MRIs acquired at the Children’s Hospital of Philadelphia (CHOP) as part of the Scans with Limited Imaging Pathology (SLIP) cohort (n=9977 T1 MPRAGE scans from 8281 participants (4259 female, 4022 male), age range 0.6–33.6 years; year of acquisition 1994–2024). SLIP scans were selected through machine-assisted manual review of radiology reports to exclude imaging pathology, as reported previously^28,29^. The secondary use of clinical scans received a waiver of consent and was determined to be exempt from further review by the CHOP Institutional Review Board because it consisted of secondary analyses of preexisting and deidentified clinical data.

### Lifespan Brain Chart Consortium datasets

The Lifespan Brain Chart Consortium (LBCC) consisted of shared and public datasets aggregated to accelerate population-scale modeling of brain development and aging. The present study included 26 LBCC datasets (n=9276 structural T1-weighted MRI scans from 6330 participants (2822 female, 3508 male), age range 0.7–88.8 years). For case-control studies, only the controls as identified by the primary study were used in the present analyses. LBCC studies, as well as the HEALthy Brain and Child Development (HBCD), Developing Human Connectome Project (dHCP) and Baby Connectome Project (BCP), are described in previous publications^12,30,31^. HBCD and dHCP also included clinical tape measures of the same participants.

### Neurogenetic cohorts

*Neurofibromatosis Type 1 (NF1):* The NF1 dataset consisted of clinically acquired scans obtained by the CHOP neurofibromatosis center as part of clinical care (n=444 participants (225 male, 219 female); n= 2071 scan sessions; Age range 1.0–21.3 years).

*22q11.2 deletion syndrome (22qDS):* The 22qDS dataset consisted of clinically acquired scans obtained by the CHOP 22q and You Center as previously described: (n=180 participants (100 male, 80 female); n= 198 scan sessions; Age range 1.8–33.5 years).

*16p11.2 deletion and duplication syndromes:* As previously described^32^, the 16p dataset consisted of 73 16p11.2 deletion carriers (41 male, 32 female; age range 1.8–49.2 years), 75

16p11.2 duplication carriers (42 male, 33 female; age range 2.1–64.3 years), 0 unaffected family members, and 0 controls.

### Cross-modal datasets

#### The CERMEP-IDB-MRXFDG (CERMEP)

As previously described^33^, the CERMEP dataset consisted of paired, same-day T1-weighted MRI and low-dose CT from 37 healthy adults (17 males and 20 females; age range, 23–65 years). T1-weighted magnetization-prepared rapid gradient-echo images were acquired at 1.5 T, and CT images were acquired during PET/CT.

#### SynthRAD2023 Grand Challenge (SynthRAD)

As previously described^34^, we used paired brain MRI (T1 weighted) and CT from the Task 1 training set of the SynthRAD2023 Grand Challenge. The training set contained 180 participants undergoing radiotherapy planning across three Dutch medical centers. Sex and training-set-specific age information were unavailable in the released metadata (full dataset reported the following for Task 1 Brain: 43.9% male, 56.1% female; age range 3–93).

### Image preprocessing and template-based registration

Each scan was first brought into a common anatomical reference space by age-conditioned template registration so that subsequent measurements were comparable across subjects and ages. Input MRI scans in NIfTI format, curated according to the brain imaging data structure (BIDS)^35^, were first registered to an age-appropriate T1-weighted template. Template selection was conditional based on the subject’s age: specific neonatal templates were used for gestational ages 36 through 44 weeks, dedicated monthly templates for ages 0 through 24 months, and broader age-range templates (e.g., NIHPD_asym_04.5-08.5_t1w) for subjects older than 2 years. The registration was performed using a rigid transformation implemented in the ANTsPy library^36^.

Axial slices were indexed along the superior–inferior axis in template space. For volumetric analyses, we processed all slices from index 40 (roughly corresponding to the chin in most participants) to the most superior slice. To mitigate residual variation in head pitch after template registration, we analyzed each volume under three rotations (0, +5, -5) about the left–right (x-) axis using a 3D Euler transform^37^. To reduce residual intensity noise before any boundary was extracted, we smoothed each slice – at every slice index and each head orientation – with curvature anisotropic diffusion (conductance parameter 3.0, 5 iterations, time step 0.0625) implemented in SimpleITK^37^.

### Binary segmentation and topology refinement

Head tissue was then separated from the image background using a combination of region-growing and intensity thresholding. First, we suppressed background voxels via connected region growing from the four image corners to a low intensity percentile, with voxels in this background region set to the minimum image intensity. The resulting image was normalized in intensity and thresholded using a valley-emphasized Otsu method (an automatic histogram-based threshold that maximizes between-class variance to separate tissue from background) with a single threshold. To ensure inclusion of low-contrast scalp and soft tissue, we lowered the Otsu threshold by a fixed modality-specific offset – −0.10 for MRI and −0.08 for CT in units of within-image intensity standard deviation – so that voxels just below the automatic cutoff, where observed scalp and soft tissue typically fall, were retained.

From these intermediate masks we selected the single head silhouette least contaminated by edge-touching, non-anatomical structures. This was done by combining the Otsu mask with the inverted corner-grown background within a narrow band around the Otsu boundary, followed by binary closing and hole filling (SimpleITK) to bridge small gaps and remove small interior voids. Among the candidate masks, we selected the one contacting the fewest image borders (ties broken by larger area), after blanking a thin border margin (4 mm for CT) to remove structures running off the fields of view. Where the selected mask still reached an image edge, we retained only the connected component overlapping an eroded (8 mm) seed, discarding thin edge- attached artefacts.

Connected-component labeling was applied to the chosen mask. When the mask touched the image borders, the component overlapping an eroded central “core” was selected; otherwise, the largest component by area was retained as the head candidate. All morphological operations were parameterized in physical units (millimeters) using the in-plane voxel spacing.

### Contour extraction

The head boundary was then extracted and cleaned at a fixed physical scale, so that artefact removal was independent of image resolution. An external binary contour was extracted from the retained component. To suppress small protrusions and close narrow indentations, all morphological operations were defined in physical units (millimeters) rather than voxels: the in-plane pixel spacing (*s_x_*, *s_y_*) was used to construct an elliptical structuring element approximating a 6-mm kernel, with each dimension rounded up to the nearest odd number of pixels so that the element had a single, well-defined centre pixel. Sequential opening then closing was applied to the binary mask, preserving the overall cranial outline, and external contours were re-extracted^38^.

Finally, the boundary was smoothed in physical space to remove residual spikes while preserving true cranial shape. To further regularize the outline while preserving global shape, the largest contour was converted to a polygon and processed in physical space^39^. First, a two-step buffer operation – erosion by 4 mm followed by dilation by 4 mm – was applied in millimeters to remove narrow spikes and bridges; if multiple polygons resulted, the largest by area was retained. The outer ring was uniformly resampled at 200 equidistant arc-length points. Coordinates were recentered to the polygon centroid and expressed as polar radii r($). Radial profiles underwent median filtering over ∼3% of the perimeter (minimum window 5) followed by Savitzky–Golay smoothing over ∼7% (window ≥11, polynomial order 2, circular “wrap” mode)^40^.

To cap residual outward spikes, each radius was limited by the local 90th percentile computed over ∼8% of the perimeter. The final smoothed contour was reconstructed from the capped radii and unit direction vectors as *c* + *r_i_u_i_*, where *u_i_* are unit radial directions.

### Circumference, shape, and asymmetry measurements

From the regularized contour and the head mask we derived head circumference together with a panel of cranial shape and asymmetry indices. Head circumference was computed as the polyline length of the smoothed contour in physical units using the in-plane spacings (*s_x_*, *s_y_*): for successive vertices, the total perimeter was calculated as 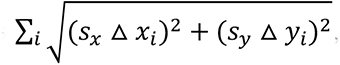, with closure enforced by connecting the last to the first vertex.

From the smoothed contour and the largest component mask, several measurements were calculated using image spacing information:

| Metric | Description | Formula |
| --- | --- | --- |
| Circumference (mm) | Perimeter of the smoothed contour. | $\sum_i \sqrt{(s_x \Delta x_i)^2 + (s_y \Delta y_i)^2}$ |
| Area (cm<sup>2</sup>) | Total area enclosed by the contour. | — |
| Quadrant Areas (cm<sup>2</sup>) | Area divided into four quadrants (posterior-left, posterior-right, anterior-left, anterior-right) based on the geometric center of the contour's bounding box; area computed for each. | — |
| Linear Dimensions (mm) | Width and length (anteroposterior diameter) from the bounding box. | — |
| Diagonal Diameters (mm) | Two diagonal diameters measured at 60° and 120° relative to the horizontal axis through the contour's centroid. | — |
| Cephalic Index (CI) | Ratio of width to length, expressed as a percentage. | $(\text{Width} / \text{Length}) \times 100$ |
| Cranial Vault Asymmetry Index (CVAI) | Asymmetry between the two diagonal diameters, expressed as a percentage. | $((\text{Longer Diagonal} - \text{Shorter Diagonal}) / \text{Longer Diagonal}) \times 100$ |
| Asymmetry Coefficient (AC) | Difference between summed left- and right-hemisphere areas, normalized by total area. | $((\text{AL} + \text{PL}) - (\text{AR} + \text{PR})) / \text{total area}$ |
| Anterior Hemisphere Asymmetry (AHA) | Left–right difference within the anterior hemisphere, normalized by anterior area. | $(\text{AL} - \text{AR}) / (\text{AL} + \text{AR})$ |
| Posterior Hemisphere Asymmetry (PHA) | Left–right difference within the posterior hemisphere, normalized by posterior area. | $(\text{PL} - \text{PR}) / (\text{PL} + \text{PR})$ |
| Cranial Roundness Index (CRI) | Overall roundness, comparing area to squared circumference; higher = more circular. | $(4 \times \pi \times \text{total\_area}) / \text{circumference\_mm}^2$ |
| Left Hemisphere Ratio (LHR) | Total left vs. right hemisphere area; > 1 indicates leftward dominance. | $(AL + PL) / (AR + PR)$ |
| Anterior–Posterior Ratio (APR) | Anterior vs. posterior area balance; larger = more frontally biased. | $(AL + AR) / (PL + PR)$ |

### Slice contour quality control metrics

Contours were represented as ordered polygon vertices on the native pixel grid (sub-pixel coordinates from the regularized outline; see Image Processing).

As quality control (QC) steps to ensure a contour was correctly processed, we produced metrics in the following categories: Convexity deficits (1), Curvature profile (2), Left–right symmetry (3).

1. *QC of* convexity deficits

We formed the convex hull (OpenCV convexHull^38^) and computed: (i) dent area = hull area − contour area (non-negative); (ii) dent ratio = dent area / hull area; and (iii) maximum hull deviation, defined as the largest inward distance from any contour vertex to the hull (via pointPolygonTest). These quantify concavities inconsistent with smooth cranial outlines^39^.

1. QC of curvature profile

We computed tangent angles along the polyline, unwrapped angle differences, and normalized by segment length to obtain discrete curvature. To suppress pixel-scale noise, curvature was smoothed using a Savitzky–Golay filter (window 9, polynomial order 3; SciPy^41^). We then summarized (i) 95th percentile of |curvature|; (ii) s.d. of |curvature|; (iii) curvature energy (integral of squared curvature); and (iv) concave fraction, the proportion of points with curvature below −0.2 (rad·px⁻¹), which flags deep inward turns.

1. QC of left–right symmetry

We measured a Hausdorff symmetry index by reflecting the contour across the vertical axis through its centroid and taking the maximum of the two directed Hausdorff distances between original and mirrored sets^41^. This penalizes asymmetric shapes typical of partial skull capture or residual extracranial tissue. A contour passed QC only if all pre-specified thresholds were simultaneously satisfied: maximum hull deviation ≤ 3.0 px; dent area ≤ 100 (px²); dent ratio ≤ 0.0045; Hausdorff symmetry ≤ 10.0 px; curvature 95th percentile ≤ 0.04 px⁻¹. All metrics were computed in pixel units unless otherwise noted, and circumference itself was always evaluated in millimeters using image spacing.

### Slice selection

The pipeline considers and evaluates all slices from index 40 in the axial plane and continues until reaching the end slice. Slice-wise area measurements were used to compute volumetric estimates for each quadrant and the total intracranial volume by multiplying the area of each slice by the slice thickness (spacing along the inferior–superior axis) and summing these contributions.

To identify the anatomically correct slice from which to extract the head circumference measurement, a secondary QC module was applied. After each slice-wise segmentation, we applied the automated QC gate to ensure geometric plausibility before accepting a contour for measurement. For each slice and tilt pair we recorded whether the measurement passed the geometric QC threshold. The pass rate for each slice was recorded as the number of tilts passing QC/ number of tilts for the given slice. The per-slice pass rate was then smoothed with a Gaussian kernel along the complete set of integer axial indices (kernel s.d. = 3 indices).

To filter scans where the output from the pipeline could be unreliable a first requirement was that the smoothed pass rate probability mass covered 40% of the axial grid indices.

Furthermore, a characteristic of a reliable output is that any slice below the axial plane of the eyes failed the geometric plausibility QC gate and everything above the eyes passed.

Therefore, a second requirement for the scan to be deemed reliable was that only one continuous band of smoothed probability mass exists for a particular scan. Any continuous probability mass covering over 10 per cent of the total integer axial indices was deemed a band; if there was more than one band, the second requirement of a reliable scan failed.

The final selected slice was the slice with maximum head circumference among the geometric QC-passing slices within the singular band.

### Manual validation of quality control procedures

To validate automated QC performance and characterize failure modes, we manually inspected 1,000 processed scans from the CHOP SLIP cohort. For each scan we generated axial slice images with overlaid cranial contours at the automatically selected measurement level, comprising the anatomical MRI slice, the segmented binary mask, and the regularized contour overlay. One rater, blinded to clinical metadata, reviewed each output in randomized order using a custom tool (Supplementary Methods) and assigned a four-point ordinal rating that distinguished contour accuracy from slice-selection accuracy. Rating 1 denoted visually acceptable contours and anatomically appropriate slice selection in all three orientations. Rating 2 denoted visually acceptable contours in all three orientations, with an anatomically appropriate slice selected in at least one orientation. Rating 3 denoted a visible contour failure in at least one orientation that automated QC successfully excluded from the final measurement. Rating 4 denoted a slice-selection failure, typically caused by registration or orientation error, despite visually acceptable contours. Ratings 1–3 were treated as scan-level passes because the final measurement was obtained from an acceptable contour and slice; rating 4 was treated as a scan-level failure. The review tool and full rubric definitions are described in the Supplementary Methods.

### Statistical analysis and centile curve modelling

All statistical analyses and visualizations were conducted using R (4.3.3). We developed sex-specific centile curves for head circumference against age using the Generalized Additive Models for Location, Scale, and Shape (GAMLSS) framework, implemented in the *gamlss* package^17^. This method allows for flexible modeling of the median, scale, skewness, and kurtosis of the response variable (head circumference) as non-linear functions of explanatory variables. Models were fit separately for females and males to allow sex-specific growth dynamics. The workflow comprised (i) age transformation; (ii) outlier screening; (iii) reference-model fitting and diagnostics; and (iv) analytic computation of centiles and z-scores.

i. *Age transformation*: Age (in days, corrected for prematurity where gestational age was available) was transformed to its base-10 logarithm to accentuate the rapid growth of early life.
ii. *Age restriction and outlier screening*: To limit the influence of extreme values before reference fitting, we fitted a preliminary regression of head circumference on age – a degree-4 B-spline of log₁₀(age) with an additive sex term – scaled the residuals by their median absolute deviation (MAD), and excluded observations whose absolute MAD-scaled residual exceeded 3.5. After outlier screening the final sample consisted of 9,685 unique participants, with one scan included per participant; 8,968 were within the range from birth to age 20. To reduce upper-boundary effects, observations beyond age 20 and up to age 41 were retained during model fitting, and the resulting curves were truncated at age 20^42,43^.
iii. *Reference-model fitting and diagnostics*: Reference curves were fitted separately for females and males using the Box–Cox power exponential (BCPE) distribution. The median (*μ*) and scale (*σ*) were each modelled as a penalized B-spline of log₁₀(age), with the effective degrees of freedom of each smoother selected by minimizing the generalized Akaike information criterion (GAIC) under a penalty of k = log(n), equivalent to the Bayesian information criterion (BIC). The skewness parameter (*v*) was held constant. The kurtosis parameter (τ) was modelled as a penalized spline of log₁₀(age) in males, where residual kurtosis was evident in the worm plots, and held constant in females. Adequacy of fit was assessed visually using worm plots (Fig S2).
iv. *Centiles and z-scores*: Centiles and z-scores were computed analytically from the fitted distribution parameters. Centile curves were obtained from the BCPE quantile function evaluated across age, qbcpe(p; *μ̂*, *σ̃*, *ν̂*, *τ̂*); each measurement’s z-score was its value mapped through the fitted BCPE cumulative distribution function and the standard-normal inverse, z = *Φ*^−1^(*F*bcpe(y; *μ̂*, *σ̃*, *ν̂*, *τ̂*)). The smooth centile curves in the figures were drawn by evaluating the parameters at the ages present in the reference sample.

Reference models for the head-shape metrics followed the same GAMLSS approach as head circumference, with the distribution family chosen to match each metric’s support and fitted separately by sex. The strictly non-negative metrics – Cephalic Index (CI), Cranial Vault Asymmetry Index (CVAI), Anterior–Posterior Ratio (APR), Cranial Roundness Index (CRI), and Left Hemisphere Ratio (LHR) – were modeled with the Box–Cox power exponential (BCPE) distribution, with the median (*μ*) and scale (*σ*) as penalized B-splines of log₁₀(age) (smoothing degrees of freedom selected by GAIC) and the skewness (*v*) and kurtosis (*τ*) held constant.

Because BCPE requires a strictly positive outcome, a small floor (ε = 10⁻³) was applied to these metrics where zero was attainable.

The signed asymmetry metrics – Asymmetry Coefficient (AC), Anterior Hemisphere Asymmetry (AHA), and Posterior Hemisphere Asymmetry (PHA) – take both positive and negative values (left- versus right-dominant) and were therefore modeled directly, without a positivity transform, using the sinh–arcsinh (SHASH) distribution, with all four parameters (*μ*, *σ*, *v*, *τ*) as penalized B-splines of log₁₀(age). As for head circumference, observations were screened for outliers by a MAD-scaled residual cut-off (|z| < 3.5), centiles and z-scores were computed analytically from the fitted parameters, and fit adequacy was assessed with worm plots. These models together with CVAI were fitted using a heavier GAIC penalty (k = 10 log n) to ensure anatomically plausible models.

### Cephalic-Group Association Models

To test whether conventional cephalic-index categories were associated with other cranial morphology features, we classified scans as dolichocephalic (CI < 75), mesocephalic (75 ≤ CI < 80), or brachycephalic (CI ≥ 80). Each cranial metric was first converted to an age- and sex-referenced z-score using the corresponding GAMLSS reference model. We then fit one linear mixed-effects model per metric, treating the metric z-score as the outcome and CI group as a categorical predictor, with mesocephalic scans as the reference group and study included as a random intercept. For non-HC metrics, HC-z was included as a covariate to estimate CI-group differences after accounting for population head-circumference deviation:

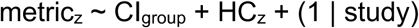

For HC itself, the model omitted HC-z: HCz ∼ CIgroup + (1 | study)

We also fit secondary interaction models for each non-HC metric to test whether the association between HC-z and each morphology metric differed by CI group: metricz ∼ CIgroup * HCz + (1 | study). P-values for all fixed effects were obtained from the Satterthwaite approximation. To account for multiple testing, all p-values across the 37 model terms tested here (dolicho- and brachy-vs meso contrasts, the HC-z covariate association, and both HC-z interaction terms, across all metrics) were corrected together as a single family using the Benjamini-Hochberg false discovery-rate procedure; significance was assessed at FDR-corrected p < 0.05.

HC-z coefficients from the additive models were interpreted as covariate associations with cranial morphology. The secondary interaction models tested whether those HC-z associations differed by CI group.

### Head-circumference-conditioned cranial-shape differences in neurogenetic cohorts

We tested for cranial-shape deviations in four neurogenetic cohorts using one scan per participant. After restricting the analysis to participants with recorded sex, an adjusted age of more than 0 and no more than 15,000 days, a valid head-circumference measurement, and finite reference z scores, the analysis included 414 participants with NF1, 63 with 16p11.2 deletion syndrome, 35 with 16p11.2 duplication syndrome, and 108 with 22q11.2 deletion syndrome. We converted head circumference and each of the eight cranial-shape measures to age- and sex-referenced z scores using the corresponding sex-specific GAMLSS reference models described above. To distinguish shape deviations from the expected association between cranial shape and global head size, we estimated a separate linear relation between each shape z score and head-circumference z score in the reference sample:

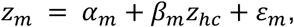

where m denotes a shape metric. For each clinical participant, we subtracted the shape z score predicted at that participant’s head-circumference z score and divided the residual by the residual standard deviation in the reference sample:

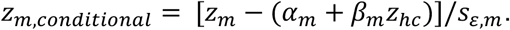

A conditional z score of zero therefore represents the shape expected given age, sex, and head-circumference deviation, and one unit represents one residual standard deviation in the reference sample. Head circumference itself remained on its marginal age- and sex-referenced z-score scale. For each cohort and metric, we tested the mean z score against zero using a two-sided one-sample t test and calculated a 95% t-based confidence interval. We controlled the false-discovery rate using the Benjamini–Hochberg procedure across all 36 comparisons (four cohorts × nine metrics, comprising head circumference and eight shape measures).

### LMSz

Each individual head circumference measure was converted to a centile by evaluating its cumulative probability under the sex-specific BCPE reference *p* = *pBCPE*(*Head circumference* | *μ̂*, *σ̂*, *ν̂*, *τ̂*) and subsequently transformed into a centile z-score by the quantile function (inverse of the cumulative distribution function) *z* = *Φ*^−1^(*p*). For measurements within the reference sample, the centile z-scores are expected to follow an approximate normal distribution *N*(0,1) across age.

To capture disease specific departure from the reference sample, we adapted the LMSz approach from which it is possible to create disorder-specific growth charts^21^. Centile z-scores for these cohorts were first produced using the reference model as described above. These z-scores from out-of-sample disease groups may have different means and variances compared to the reference sample. The out-of-sample z-scores were thus modelled by fitting a secondary GAMLSS model using the normal distribution family where candidate formulas were again tested by grid search and evaluated through BIC scores.

To get the LMSz z-score for the cohort we use the secondary GAMLSS model to create cohort-specific LMSz z-scores as *z_c_*(*age*, *sex*; *p*) = *μ_z_*(*age*, *sex*) + *σ*(*age*, *sex*)*z_p_*, for percentile *p* and *z_p_* = *Φ*^−1^(*p*). This cohort-specific z-score is back-transformed into the units of the original measurements by a simple p-norm transformation of the z-score followed by *qBCPE*(*p_c_ μ̂*, *σ̂*, *ν̂*, *τ̂*). Cohort-specific centile lines were plotted for *p* = {3, 50, 97}.

### Correlations with tape measurements and replicability of measurements

To validate the imaging-derived head circumference against the clinical reference standard, we compared it with tape-measured head circumference in four cohorts (HBCD, 16p, SLIP2025, and dHCP). Each MRI scan was paired with the closest available tape measurement: in HBCD and dHCP, the tape head circumference recorded in the same session as the scan; in 16p, the tape measurement at the same age in months (the scan’s chronological age rounded to the nearest month) for that individual; and in SLIP2025, where no paired research measurement existed, the nearest routine clinical head-circumference measurement within 30 days (restricted to physiologically plausible values, 30–60 cm). Where multiple same-session tape readings were available (HBCD), they were collapsed to a single value by averaging concordant readings (within 0.2 cm) and otherwise taking the median.

Agreement was assessed by Bland–Altman analysis: for each cohort we computed the bias (mean of MRI minus tape) and the 95% limits of agreement (bias ± 1.96 × s.d. of the differences), expressed both in raw centimeters and in age- and sex-adjusted z-score units. Tape measurements were converted to z-scores using the CDC 2000 and UK-WHO growth references and MRI measurements using the BCPE reference models described above^1,18–20^; outlying pairs were removed per cohort before analysis.

Measurement replicability was quantified with one-way random-effects single-measure intraclass correlation coefficients (ICC(1,1), Shrout–Fleiss notation), treating the repeated measurements within a session as exchangeable replicates. We computed this for MRI repeatability, across the repeat scans acquired within a single imaging session.

To characterize the variability inherent to the reference method, we also quantified the test– retest consistency of the physical tape measurements in the HBCD cohort, where repeated same-session readings were available (622 sessions with three readings – two taken on the same occasion by one examiner and a third later the same day by a second examiner – and 14 sessions with two readings by a single examiner). Each reading was converted to a z-score using the CDC 2000 and UK-WHO LMS references, and within-session consistency was summarized as a mean absolute error (MAE): the mean absolute deviation of the repeated readings from their session mean, computed per session and averaged across sessions, on the CDC-z, UK-WHO-z, and raw-cm scales.

### Disclosures

AA-B, AZ, RAIB, JDB, and JS hold shares in, AA-B and PM has consulted for, and JDB and JS are directors of Centile Bioscience. PM, MG, BJ, SK, DZ, EL, AA-B, and JS have an inventorship interest in CHOP IP licensed to Centile Bioscience. A provisional patent has been filed related to this work. JDB also has an equity position in Treovir Inc., is the CMO of UpFront Diagnostics, is a co-founder of Oncolight, and is on the QV Bioelectronics and Quadrillion boards of scientific advisors. TR declares consulting or advisory board relationships and/or equity positions in Prism Clinical Imaging, Proteus Neurodynamics, Fieldline Medical Inc. and WestCan Proton Therapy Inc. RTS has received consulting income from Octave Bioscience and Sanofi.

## Acknowledgements

This work was funded by NIMH R01MH134896, NIMH R01MH133843, and the CHOP Research Institute. G.B was supported by an NHMRC Investigator Grant (1194497), JAB is supported by the *Eunice Kennedy Shriver* National Institute of Child Health and Human Development through the Pediatric Research Loan Repayment Program (L40HD119861).

## Correspondence

Jakob Seidlitz or Aaron Alexander- Bloch: Richards Medical Research Laboratories, 3700 Hamilton Walk, Philadelphia, PA 19104

