## Supplement for "Charting neuroimaging-based head circumference and cranial shape across development"

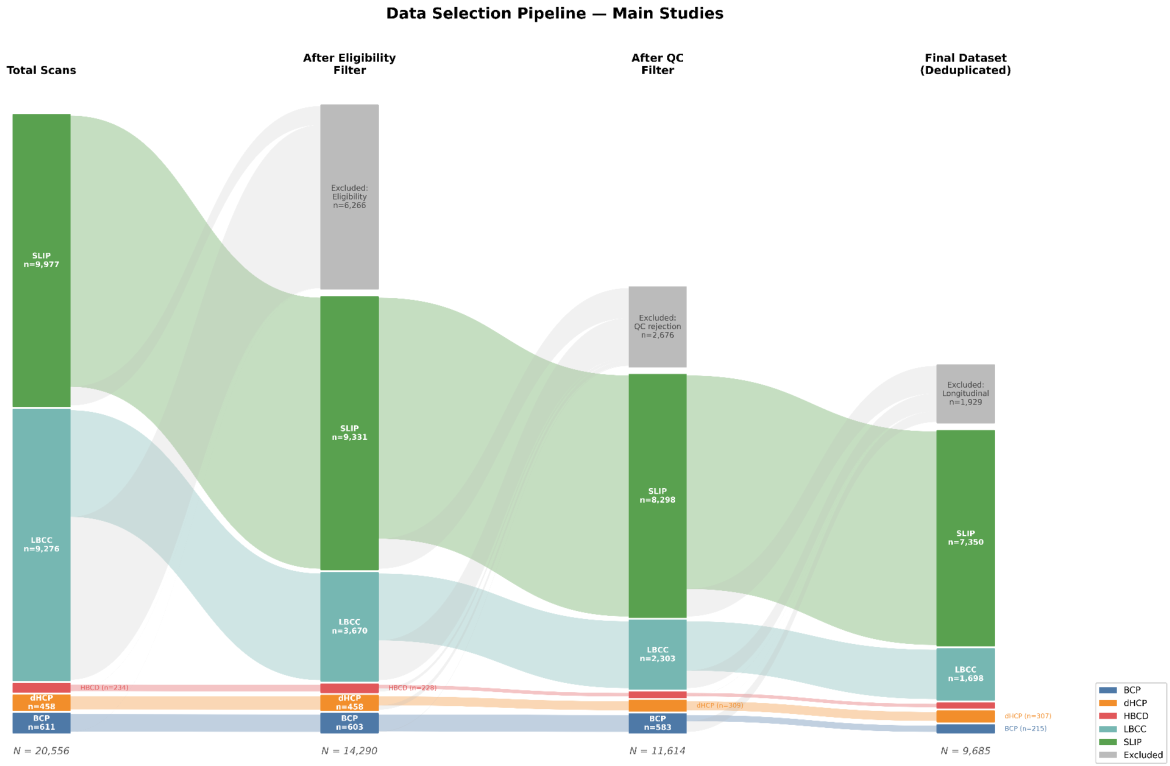


**Figure S1**. Data-selection pipeline for the population reference sample. Sankey diagram tracking scans from the full pool to the final deduplicated analytic sample (N = 9,685), by study (SLIP, LBCC, dHCP, BCP, HBCD). Scans were removed at three stages: eligibility filtering, automated quality-control rejection, and de-duplication of longitudinal or repeat acquisitions. The final sample comprised SLIP (7,350), LBCC (1,698), dHCP (307), BCP (215), and HBCD (115) scans. Column totals are shown beneath each stage.

### Supplemental Methods

#### *Manual quality-control review*

We generated a custom Python-based graphical user interface that displayed each image sequentially and prompted for rating input via keyboard shortcuts (keys 1-4) or mouse click. The interface positioned a compact rating dialog in the top-right corner of the screen, allowing simultaneous viewing of the full anatomical image and rating controls without window switching. The dialog remained topmost to prevent occlusion by the image viewer window. Images were presented in randomized order to prevent order effects and raters were blinded to clinical metadata. All ratings were timestamped and recorded to a comma-separated values (CSV) file containing image filename, rating value (1-4), rater identifier, and timestamp.

One rater independently reviewed each output using a standardized four-point ordinal scale designed to distinguish contour accuracy from slice-selection accuracy. Ratings were based primarily on visual inspection of the contours and selected slices. Automated QC status was considered only when determining whether a visibly defective orientation had been excluded from the final measurement.

Rating 1 (Complete pass): The external head contour was visually accurate, the selected slice was anatomically appropriate – that is, at or above the orbits and capturing the maximum circumference – and all three orientations yielded acceptable outputs.

Rating 2 (Pass with partial slice-selection agreement): Contours were visually acceptable in all three orientations, but an anatomically appropriate slice was selected in only one or two orientations.

Rating 3 (Contour failure excluded): At least one orientation showed a visible contour failure, such as excessive concavity, marked asymmetry, field-of-view truncation, or tracing of an internal rather than external boundary. Automated QC excluded the defective orientation, and the final measurement was obtained from an acceptable contour and slice in another orientation.

Rating 4 (Slice-selection failure): Contours were visually acceptable, but the pipeline selected an anatomically incorrect slice because of registration or orientation error. The selected slice was too superior, too inferior, or included orbital or facial structures.

Ratings 1–3 were treated as scan-level passes because the final measurement was obtained from an acceptable contour and slice. Rating 4 was treated as a scan-level failure. This rating scheme distinguished orientation-level contour failures successfully excluded by automated QC from slice-selection failures that affected the final measurement.

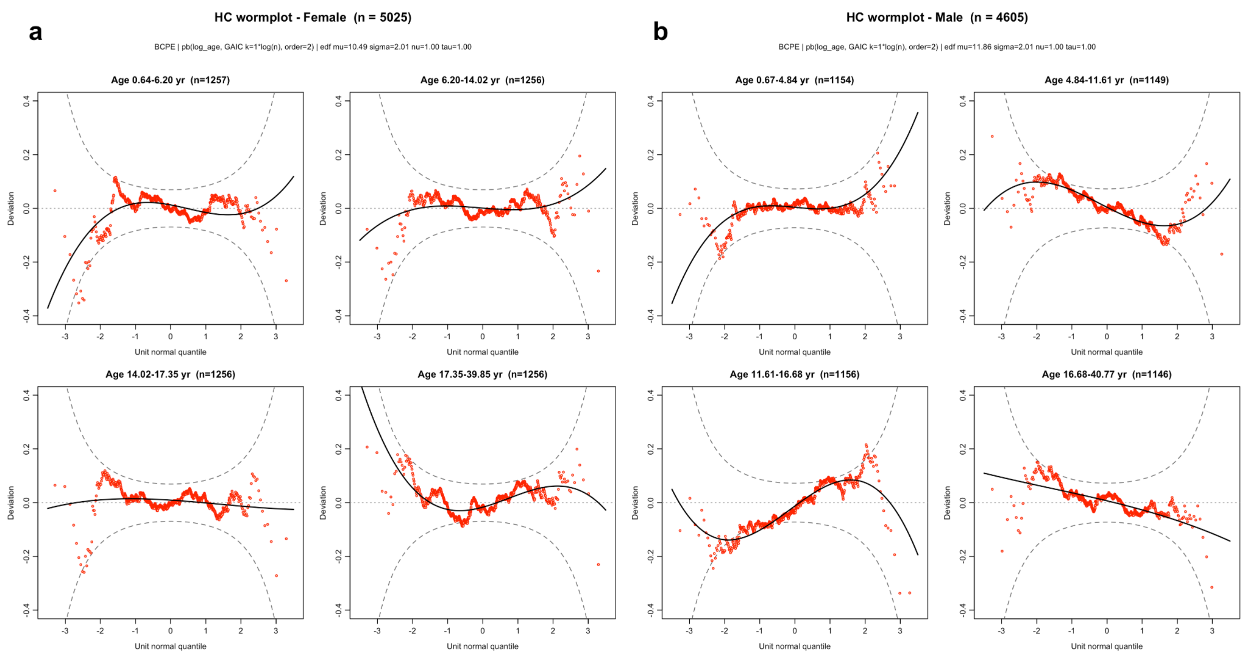


**Figure S2.** Worm-plot diagnostics for the head-circumference reference model. Worm plots (detrended normal quantile–quantile plots of the normalized quantile residuals) for the sex-specific BCPE head-circumference model, in (a) females and (b) males, faceted into four equal-count age bins (years; bin limits shown in each panel header). Points are the residuals, the solid curve is a cubic trend, and the dashed curves are the pointwise 95% envelope. Residuals scattered about zero and lying within the envelope indicate adequate calibration across age.

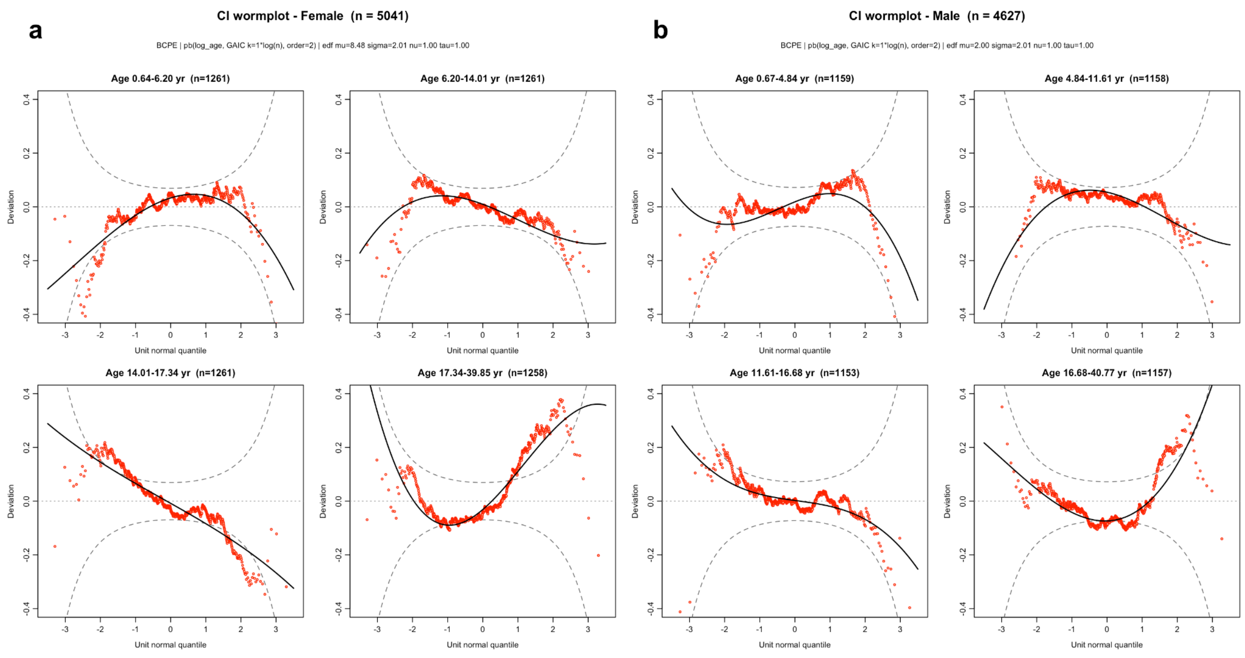


**Figure S3.** Worm-plot diagnostics for the Cephalic Index (CI) reference model. Worm plots for the sex-specific BCPE CI model, in (a) females (n = 5,041) and (b) males (n = 4,627), faceted into four equal-count age bins (years; bin limits in each panel header). Red points are the normalized quantile residuals, the solid curve is a cubic trend, and the dashed curves are the pointwise 95% envelope; residuals near zero and within the envelope indicate adequate fit.

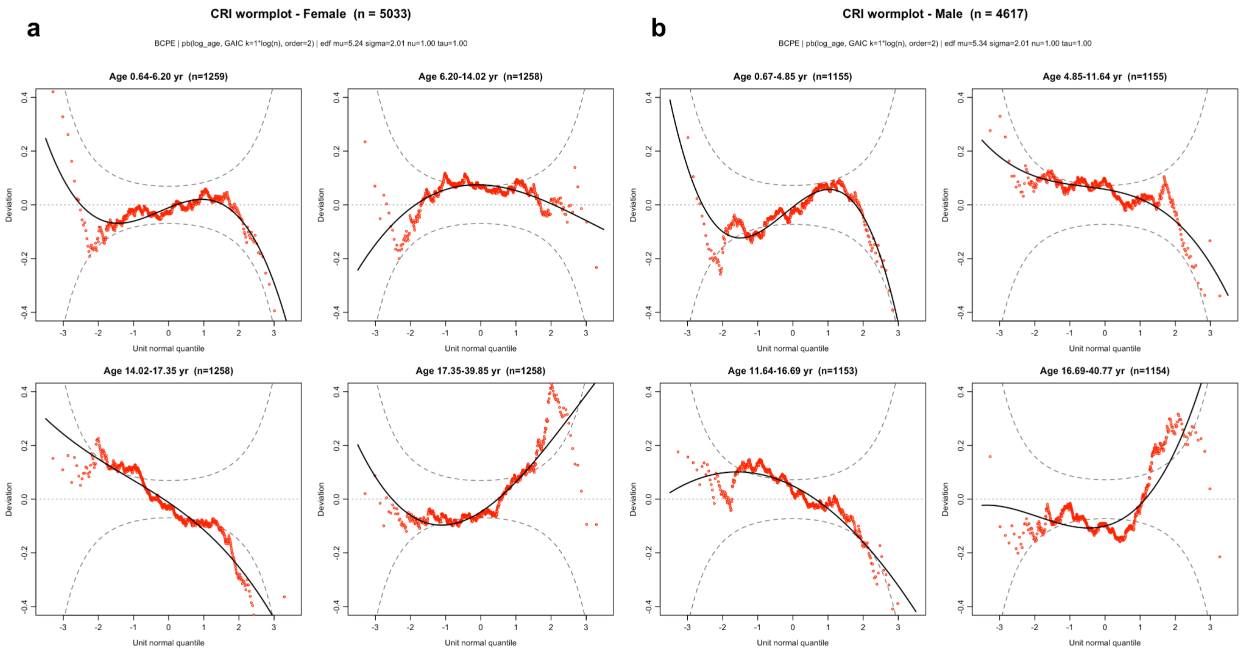


**Figure S4.** Worm-plot diagnostics for the Cranial Roundness Index (CRI) reference model. Wormplots for the sex-specific BCPE CRI model, in (a) females (n = 5,033) and (b) males (n = 4,617), faceted into four equal-count age bins (years; bin limits in each panel header). Red points are the normalized quantile residuals, the solid curve is a cubic trend, and the dashed curves are the pointwise 95% envelope; residuals near zero and within the envelope indicate adequate fit.

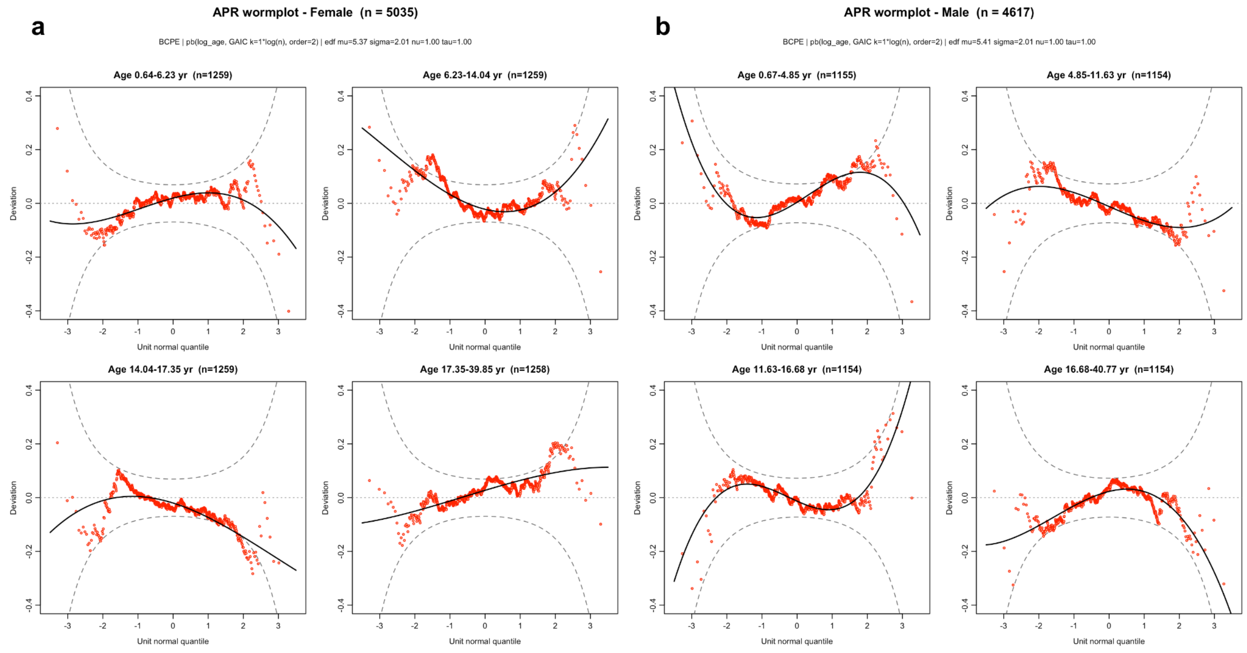


**Figure S5**. Worm-plot diagnostics for the Anterior–Posterior Ratio (APR) reference model. Worm plots for the sex-specific BCPE APR model, in (a) females (n = 5,035) and (b) males (n = 4,617), faceted into four equal-count age bins (years; bin limits in each panel header). Red points are the normalized quantile residuals, the solid curve is a cubic trend, and the dashed curves are the pointwise 95% envelope; residuals near zero and within the envelope indicate adequate fit.

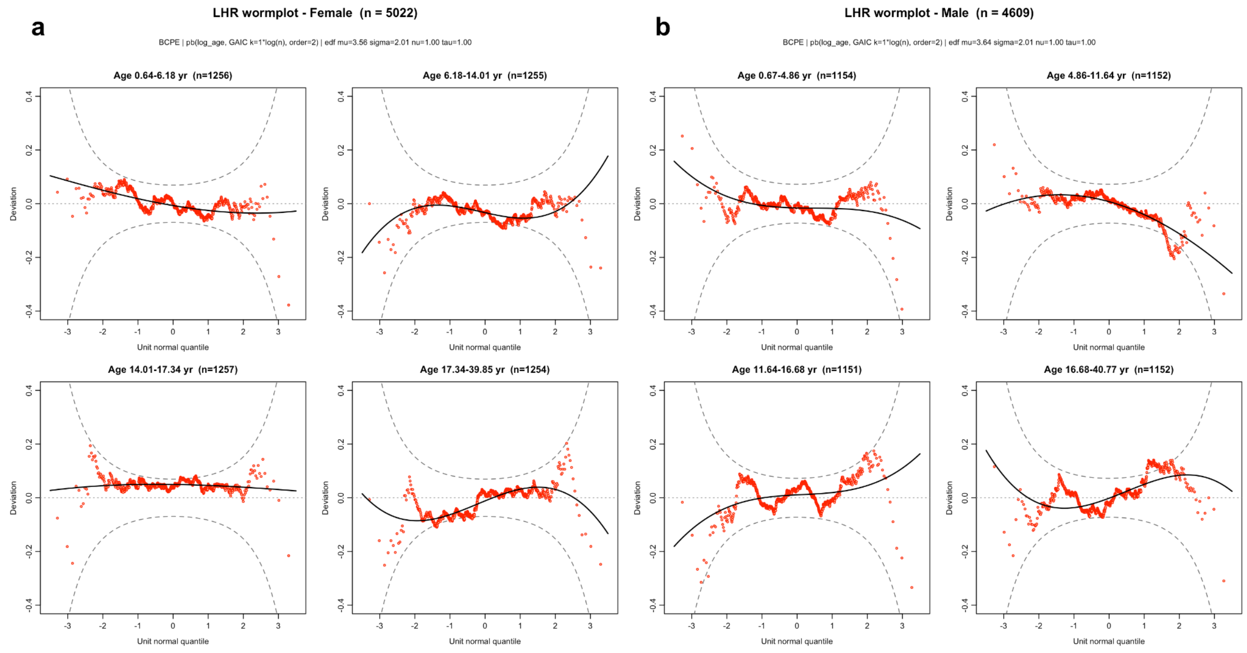


**Figure S6.** Worm-plot diagnostics for the Left Hemisphere Ratio (LHR) reference model. Worm plots for the sex-specific BCPE LHR model, in (a) females (n = 5,022) and (b) males (n = 4,609), faceted into four equal-count age bins (years; bin limits in each panel header). Red points are the normalized quantile residuals, the solid curve is a cubic trend, and the dashed curves are the pointwise 95% envelope; residuals near zero and within the envelope indicate adequate fit.

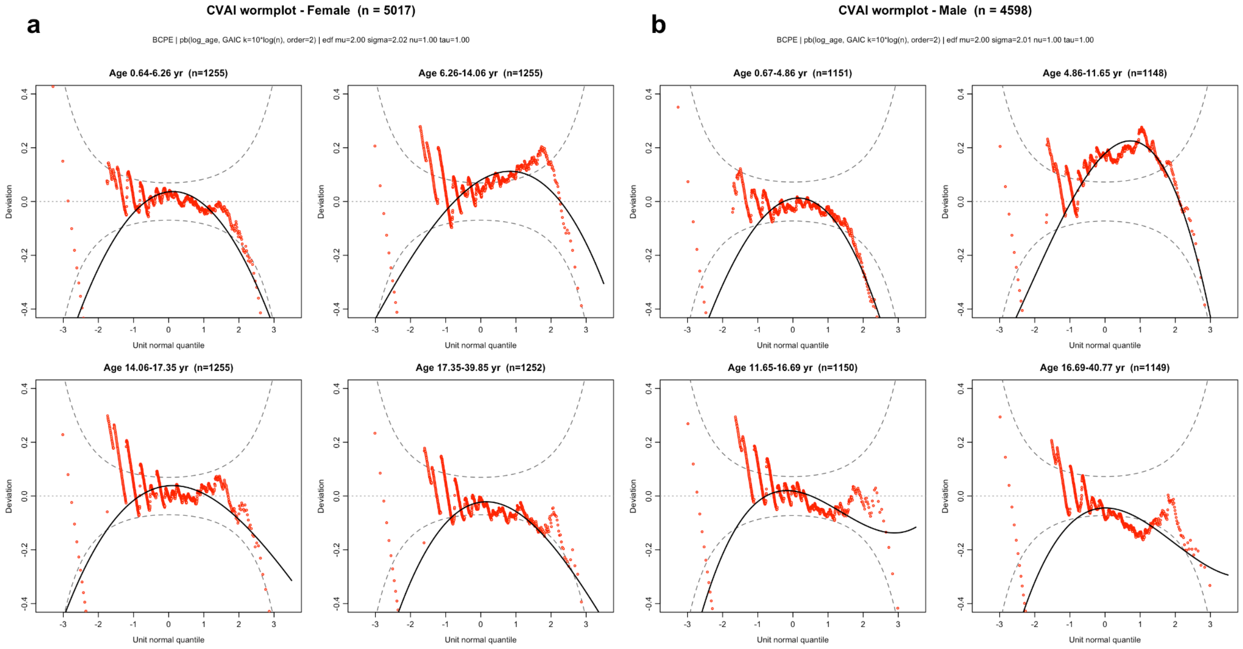


**Figure S7.** Worm-plot diagnostics for the Cranial Vault Asymmetry Index (CVAI) reference model. Worm plots for the sex-specific BCPE model of CVAI (taken as an absolute value), in (a) females (n = 5,017) and (b) males (n = 4,598), faceted into four equal-count age bins (years; bin limits in each panel header). Red points are the normalized quantile residuals, the solid curve is a cubic trend, and the dashed curves are the pointwise 95% envelope. The vertical striping at low values reflects the floor at zero inherent to the absolute-value metric.

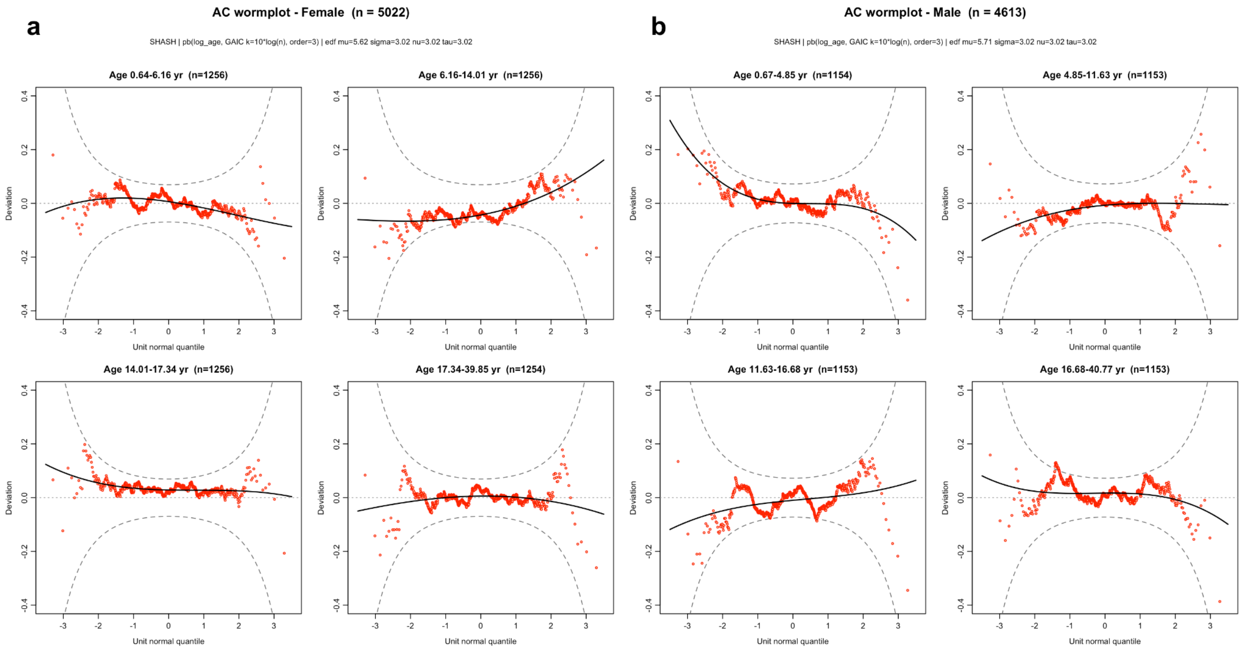


**Figure S8.** Worm-plot diagnostics for the Asymmetry Coefficient (AC) reference model. Worm plots for the sex-specific SHASH model of the signed AC, in (a) females (n = 5,022) and (b) males (n = 4,613), faceted into four equal-count age bins (years; bin limits in each panel header). Red points are the normalized quantile residuals, the solid curve is a cubic trend, and the dashed curves are the pointwise 95% envelope. AC is modeled on its signed (left-minus-right) value, so positive and negative deviations denote opposite directions of asymmetry.

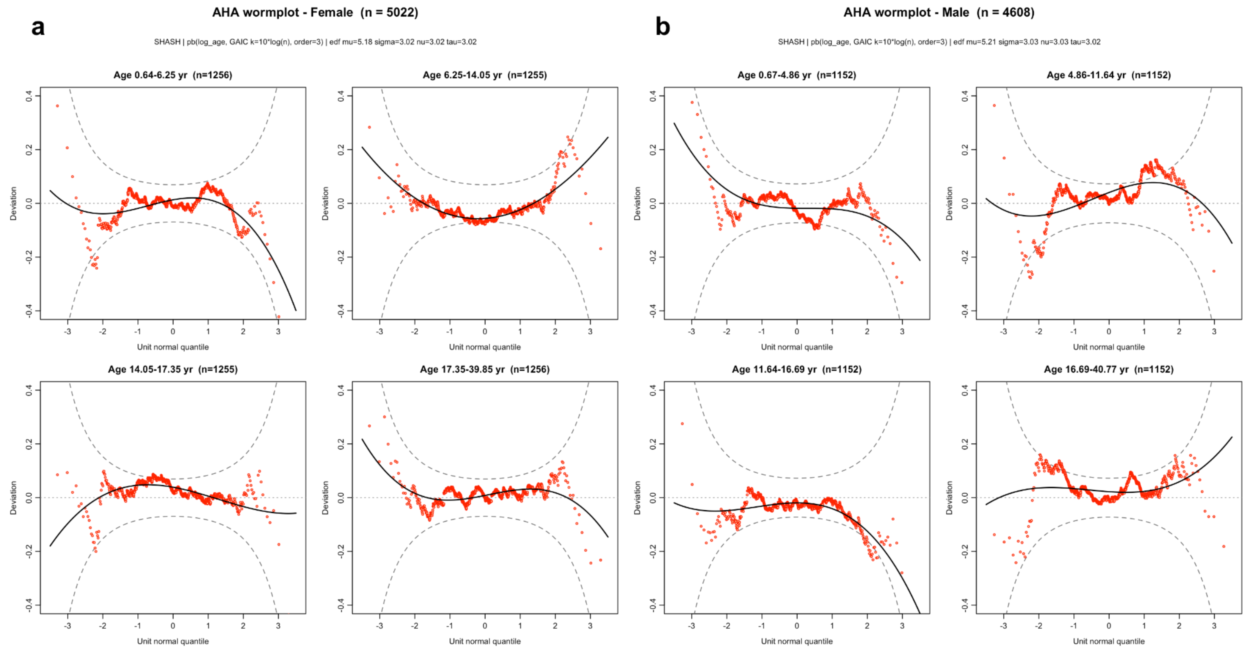


**Figure S9.** Worm-plot diagnostics for the Anterior Hemisphere Asymmetry (AHA) reference model. Worm plots for the sex-specific SHASH model of the signed AHA, in (a) females (n = 5,022) and (b) males (n = 4,608), faceted into four equal-count age bins (years; bin limits in each panel header). Red points are the normalized quantile residuals, the solid curve is a cubic trend, and the dashed curves are the pointwise 95% envelope. AHA is modeled on its signed (left-minus-right) value, so positive and negative deviations denote opposite directions of asymmetry.

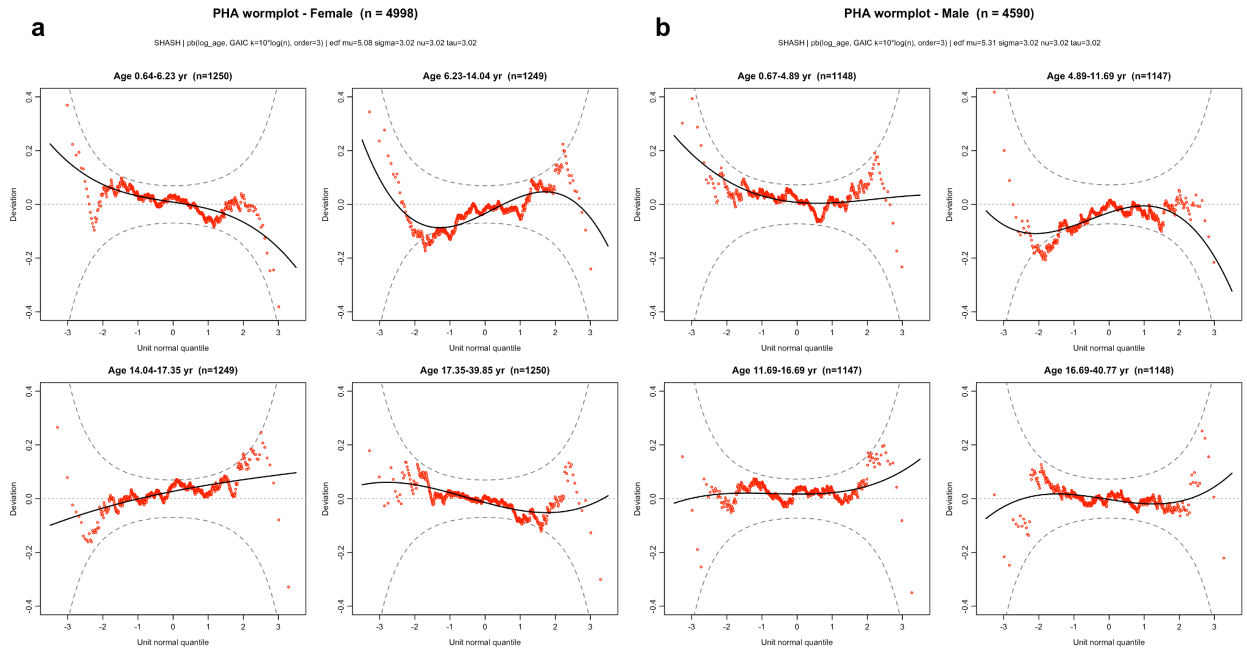


**Figure S10.** Worm-plot diagnostics for the Posterior Hemisphere Asymmetry (PHA) reference model.

Worm plots for the sex-specific SHASH model of the signed PHA, in (a) females (n = 4,998) and (b) males (n = 4,590), faceted into four equal-count age bins (years; bin limits in each panel header). Red points are the normalized quantile residuals, the solid curve is a cubic trend, and the dashed curves are the pointwise 95% envelope. PHA is modeled on its signed (left-minus-right) value; consistent with the main-text finding, posterior asymmetry is predominantly right-dominant.

**Supplementary Table 1.** Head-circumference and cranial-shape deviations in four neurogenetic cohorts. For each cohort–metric comparison, the table reports the number of independent participants, mean z score, standard deviation, t statistic, degrees of freedom, 95% confidence interval, two-sided one-sample t-test P value, and Benjamini–Hochberg-adjusted P value. Head circumference is referenced to age and sex; each shape metric is additionally conditioned on the linear association with head-circumference z score in the reference sample. Adjustment was applied across all 36 cohort–metric comparisons.

| cohort | phenotype | metric | n | mean_z | sd_z | t | df | p | se | ci_low | ci_high | adjustment | p_BH | sig |
|---|---|---|---|---|---|---|---|---|---|---|---|---|---|---|
| NF1 | macro | HC | 414 | 0.71 | 0.95 | 15.14 | 413 | < 0.001 | 0.05 | 0.62 | 0.80 | age_sex | < 0.001 | *** |
| NF1 | macro | CI | 414 | -0.12 | 0.92 | -2.60 | 413 | 0.010 | 0.05 | -0.21 | -0.03 | age_sex_HC | 0.044 | * |
| NF1 | macro | AC | 414 | 0.07 | 0.97 | 1.45 | 413 | 0.148 | 0.05 | -0.02 | 0.16 | age_sex_HC | 0.411 | |
| NF1 | macro | CVAI | 414 | 0.19 | 1.00 | 3.77 | 413 | < 0.001 | 0.05 | 0.09 | 0.28 | age_sex_HC | < 0.001 | *** |
| NF1 | macro | AHA | 414 | 0.11 | 0.99 | 2.35 | 413 | 0.019 | 0.05 | 0.02 | 0.21 | age_sex_HC | 0.077 | . |
| NF1 | macro | PHA | 414 | -0.03 | 0.97 | -0.63 | 413 | 0.530 | 0.05 | -0.12 | 0.06 | age_sex_HC | 0.764 | |
| NF1 | macro | CRI | 414 | -0.18 | 0.96 | -3.85 | 413 | < 0.001 | 0.05 | -0.27 | -0.09 | age_sex_HC | < 0.001 | *** |
| NF1 | macro | APR | 414 | 0.35 | 1.05 | 6.86 | 413 | < 0.001 | 0.05 | 0.25 | 0.46 | age_sex_HC | < 0.001 | *** |
| NF1 | macro | LHR | 414 | 0.07 | 0.97 | 1.51 | 413 | 0.132 | 0.05 | -0.02 | 0.17 | age_sex_HC | 0.395 | |

| | | | | | | | | | | | | | | |
|---|---|---|---|---|---|---|---|---|---|---|---|---|---|---|
| 16p_deletion | macro | HC | 63 | 0.60 | 0.91 | 5.22 | 62 | < 0.001 | 0.12 | 0.37 | 0.83 | age_sex | < 0.001 | *** |
| 16p_deletion | macro | CI | 63 | -0.15 | 0.82 | -1.41 | 62 | 0.162 | 0.10 | -0.35 | 0.06 | age_sex_HC | 0.417 | |
| 16p_deletion | macro | AC | 63 | -0.14 | 0.99 | -1.10 | 62 | 0.276 | 0.12 | -0.38 | 0.11 | age_sex_HC | 0.621 | |
| 16p_deletion | macro | CVAI | 63 | 0.01 | 0.97 | 0.06 | 62 | 0.951 | 0.12 | -0.24 | 0.25 | age_sex_HC | 0.978 | |
| 16p_deletion | macro | AHA | 63 | -0.11 | 0.99 | -0.86 | 62 | 0.390 | 0.12 | -0.36 | 0.14 | age_sex_HC | 0.724 | |
| 16p_deletion | macro | PHA | 63 | -0.08 | 0.90 | -0.71 | 62 | 0.481 | 0.11 | -0.31 | 0.15 | age_sex_HC | 0.724 | |
| 16p_deletion | macro | CRI | 63 | 0.00 | 0.84 | 0.01 | 62 | 0.992 | 0.11 | -0.21 | 0.21 | age_sex_HC | 0.992 | |
| 16p_deletion | macro | APR | 63 | -0.05 | 0.88 | -0.44 | 62 | 0.663 | 0.11 | -0.27 | 0.17 | age_sex_HC | 0.770 | |
| 16p_deletion | macro | LHR | 63 | -0.13 | 0.98 | -1.04 | 62 | 0.303 | 0.12 | -0.38 | 0.12 | age_sex_HC | 0.641 | |
| 16p_duplication | micro | HC | 35 | -1.07 | 1.10 | -5.77 | 34 | < 0.001 | 0.19 | -1.45 | -0.69 | age_sex | < 0.001 | *** |
| 16p_duplication | micro | CI | 35 | -0.01 | 0.89 | -0.08 | 34 | 0.936 | 0.15 | -0.32 | 0.29 | age_sex_HC | 0.978 | |
| 16p_duplication | micro | AC | 35 | -0.08 | 1.14 | -0.41 | 34 | 0.681 | 0.19 | -0.47 | 0.31 | age_sex_HC | 0.770 | |
| 16p_duplication | micro | CVAI | 35 | -0.14 | 1.14 | -0.71 | 34 | 0.483 | 0.19 | -0.53 | 0.25 | age_sex_HC | 0.724 | |
| 16p_duplication | micro | AHA | 35 | -0.22 | 1.14 | -1.14 | 34 | 0.263 | 0.19 | -0.61 | 0.17 | age_sex_HC | 0.621 | |

| | | | | | | | | | | | | | | |
|---|---|---|---|---|---|---|---|---|---|---|---|---|---|---|
| 16p_duplication | micro | PHA | 35 | 0.15 | 1.06 | 0.83 | 34 | 0.413 | 0.18 | -0.22 | 0.51 | age_sex_HC | 0.724 | |
| 16p_duplication | micro | CRI | 35 | 0.09 | 0.96 | 0.52 | 34 | 0.603 | 0.16 | -0.24 | 0.41 | age_sex_HC | 0.770 | |
| 16p_duplication | micro | APR | 35 | -0.08 | 1.01 | -0.47 | 34 | 0.640 | 0.17 | -0.43 | 0.27 | age_sex_HC | 0.770 | |
| 16p_duplication | micro | LHR | 35 | -0.07 | 1.13 | -0.38 | 34 | 0.706 | 0.19 | -0.46 | 0.32 | age_sex_HC | 0.770 | |
| 22q | micro | HC | 108 | -0.59 | 1.13 | -5.45 | 107 | < 0.001 | 0.11 | -0.81 | -0.38 | age_sex | < 0.001 | *** |
| 22q | micro | CI | 108 | 0.08 | 1.06 | 0.76 | 107 | 0.447 | 0.10 | -0.12 | 0.28 | age_sex_HC | 0.724 | |
| 22q | micro | AC | 108 | -0.04 | 1.04 | -0.44 | 107 | 0.659 | 0.10 | -0.24 | 0.15 | age_sex_HC | 0.770 | |
| 22q | micro | CVAI | 108 | -0.19 | 1.00 | -1.97 | 107 | 0.051 | 0.10 | -0.38 | 0.00 | age_sex_HC | 0.185 | |
| 22q | micro | AHA | 108 | 0.09 | 0.98 | 0.95 | 107 | 0.347 | 0.09 | -0.10 | 0.28 | age_sex_HC | 0.693 | |
| 22q | micro | PHA | 108 | -0.14 | 0.91 | -1.60 | 107 | 0.113 | 0.09 | -0.31 | 0.03 | age_sex_HC | 0.370 | |
| 22q | micro | CRI | 108 | -0.05 | 0.95 | -0.55 | 107 | 0.584 | 0.09 | -0.23 | 0.13 | age_sex_HC | 0.770 | |
| 22q | micro | APR | 108 | 0.07 | 1.03 | 0.74 | 107 | 0.463 | 0.10 | -0.12 | 0.27 | age_sex_HC | 0.724 | |
| 22q | micro | LHR | 108 | -0.04 | 1.04 | -0.39 | 107 | 0.696 | 0.10 | -0.24 | 0.16 | age_sex_HC | 0.770 | |
